# Root system influence on high dimensional leaf phenotypes over the grapevine growing season

**DOI:** 10.1101/2020.11.10.376947

**Authors:** Zachary N. Harris, Laura L. Klein, Mani Awale, Joel F. Swift, Zoë Migicovsky, Niyati Bhakta, Emma Frawley, Daniel H. Chitwood, Anne Fennell, Laszlo G. Kovacs, Misha Kwasniewski, Jason P. Londo, Qin Ma, Allison J. Miller

## Abstract

- In many perennial crops, grafting the root system of one individual to the shoot system of another individual has become an integral part of propagation performed at industrial scales to enhance pest, disease, and stress tolerance and to regulate yield and vigor. Grafted plants offer important experimental systems for understanding the extent and seasonality of root system effects on shoot system biology.
- Using an experimental vineyard where a common scion ‘Chambourcin’ is growing ungrafted and grafted to three different rootstocks, we explore associations between root system genotype and leaf phenotypes in grafted grapevines across a growing season. We quantified five high-dimensional leaf phenotyping modalities: ionomics, metabolomics, transcriptomics, morphometrics, and physiology and show that rootstock influence is subtle but ubiquitous across modalities.
- We find strong signatures of rootstock influence on the leaf ionome, with unique signatures detected at each phenological stage. Moreover, all phenotypes and patterns of phenotypic covariation were highly dynamic across the season.
- These findings expand upon previously identified patterns to suggest that the influence of root system on shoot system phenotypes is complex and broad understanding necessitates volumes of high-dimensional, multi-scale data previously unmet.

## Introduction

High-throughput data acquisition has afforded unprecedented capacity to quantify and understand plant phenotypes. Recent advances in imaging and computation have expanded our ability to measure plant form (Ubbens & Stavness, 2017; Gehan *et al*., 2017), and to extend those comprehensive measurements into latent space phenotypes (Ubbens *et al*., 2020). Phenomics is characterized as the acquisition and analysis of high-dimensional phenotypic data at hierarchical levels (Soulé, 1967; Houle *et al*., 2010), often with an eye toward multiscale data integration. This holistic and hierarchical approach to plant form and function affords unique insight into how plants change over developmental time, and in response to environmental cues and horticultural manipulation.

One common horticultural manipulation is grafting, the ancient agricultural practice that joins the stem of one plant (the scion) with the root system of another plant (the rootstock) (Mudge *et al*., 2009). In agriculture, grafting is commonly used to confer favorable phenotypes that preferred scions lack. Such phenotypes include enhanced disease resistance (Pouget, 1990; Walker *et al*., 2014), fruit quality, plant form (Warschefsky *et al*., 2016), response to water stress (Tramontini *et al*., 2013), and growth on particular soils (Bavaresco & Lovisolo, 2015; Ferlito *et al*., 2020). Because grafting involves the union of a scion with a different (genetically distinct) rootstock, it offers a valuable experimental system in which root system impacts on shoot system phenotypes can be evaluated.

The cultivated grapevine, *Vitis* spp., is among the most economically important fruit crops in the world. Grapevines are cultivated primarily for fruits used to make wine and juice, as well as for table grape and raisin production. Most work on the molecular response to grafting in grapevine shows a remarkable breadth of scion response patterns. For example, a study of ‘Cabernet Sauvignon’ grafted to different rootstocks identified transcriptome reprogramming in the scion of grafted plants; this appeared to be a general effect of grafting to a rootstock and was not rootstock-specific (Cookson & Ollat, 2013). In contrast, other studies have found signatures of rootstock genotype in the transcriptome in early berry development, although this distinction was lost in later development (Berdeja *et al*., 2015; Corso *et al*., 2016), but see (Zombardo *et al*., 2020). Collectively, these studies suggest the effects of grafting are diverse and may vary over the course of vine development.

Comprehensive phenomic analyses, including those that link transcriptome data with other high-throughput phenotypic assays, offer an opportunity to expand understanding of grafting effects on grapevine shoots. For example, leaves of the cultivar ‘Gaglioppo’ show variation in stilbene and abscisic acid concentrations due to rootstock genotype, as well as differences in transcriptional profiles (Chitarra *et al*., 2017). Likewise, gene expression, ion concentrations, and leaf shape in the cultivar ‘Chambourcin’ varied in response to rootstock genotype (Chitarra *et al*., 2017; Migicovsky *et al*., 2019a). Nonetheless, questions remain regarding variation imparted by grafting over the course of the growing season and the extent to which different phenotypes covary.

Grapevine leaves are the main photosynthetic engine of the organism and a primary site for perception and response to environmental change. Leaves present a wide variety of highly variable and readily assayable phenotypes, providing an important opportunity for comprehensive phenomic assessment. Grapevine leaves have been used for centuries as markers of species and cultivar delimitation, developmental variation, disease presence, and nutrient deficiency (Galet, 1979; Mullins *et al*., 1992). More recently, analysis of grapevine leaf morphology has identified genetic architecture of leaf shapes (Chitwood *et al*., 2014), developmental patterns across the season (Chitwood *et al*., 2015), and signatures of evolution in the grapevine genus (Klein *et al*., 2017). Grapevine leaves respond to stress through gas and water exchange with the atmosphere (Williams & Grimes, 1987; Grimes & Williams, 1990) and have been shown to differentially partition the ionome depending on their position on the shoot (Migicovsky *et al*., 2019a) and their rootstock genotype (Lecourt *et al*., 2015; Migicovsky *et al*., 2019a; Gautier *et al*., 2020a). The volume of work on grapevine leaves provides a foundation for the analysis of phenomic variation in a vineyard over a season in response to grafting.

In this study, we investigate effects of seasonal variation and grafting on leaf phenomic variation of the hybrid cultivar ‘Chambourcin’. We show that ionomic, metabolomic, transcriptomic, morphometric, and physiology phenotypes vary over the course of the season and reflect subtle but ubiquitous responses to grafting and rootstock genotype. Rootstock effects were often dynamic across the season, suggesting that accounting for seasonal variation could alter our understanding of grafting in viticulture.

## Methods

### Study Design

Data were collected in an experimental rootstock trial at the University of Missouri’s Southwest Research Center (37.074167 N; 93.879167 W). Samples were collected in 2017 at three phenological stages: anthesis (~80% of open flowers; 22 May 2017); veraison (~50% of berries had transitioned from green to red; 30 July 2017); and immediately prior to harvest (25 September 2017). The vineyard includes the interspecific hybrid cultivar ‘Chambourcin’ growing ungrafted (own-rooted) and grafted to three rootstocks: ‘1103P’, ‘3309C’, and ‘SO4’. Each of the four rootstock-scion combinations was replicated 72 times for a total of 288 vines planted in nine rows. Each row was treated with one of three irrigation treatments: full evapotranspiration replacement, partial (50%) evapotranspiration replacement (reduced deficit irrigation; RDI), or no evapotranspiration replacement. Rainfall in 2017 likely mitigated the applied irrigation treatment (see: Supplemental Note 1). Vine position in the vineyard corresponded to time of sampling for some phenotypes, as samples were taken from one end of the vineyard to the other over the course of two to three hours. Because vineyard microclimates and sampling time may be associated with phenotypic variation, we defined ‘temporal block’ as a factor that captures this spatial and temporal variation inherent in sampling. Unique rootstock-scion combinations were planted in cells of four adjacent replicated vines, with rows consisting of eight cells. Depending on the phenotype being assayed, leaves were sampled from either the full vineyard (the 288-vine set) or from a nested set comprising 72 vines representing the middle two vines in each four-vine cell (the 72-vine set).

### Leaf Ionomics

The ionome describes concentrations of ions in a tissue at a particular time point (Salt *et al*., 2008). From the 288-vine set, three leaves were collected along a single shoot: the youngest fully opened leaf at the shoot tip, the approximate middle leaf, and the oldest leaf at the shoot base. Leaves were sampled from primary shoots, placed in zip-lock bags in the field and dried in coin envelopes at 50°C for one to three days. Between 20 and 100 mg of leaf tissue was acid digested and 20 ions were quantified using inductively coupled plasma mass spectrometry (ICP-MS) following standard protocol (Baxter, 2010; Ziegler *et al*., 2013)at the Donald Danforth Plant Science Center (DDPSC). Ion quantifications were corrected for sample losses, internal standard concentrations, instrument drift and by initial sample mass as part of the DDPSC Ionomics Pipeline. For each ion concentration, we computed z-score distributions and used those values as the basis for linear models. Non-standardized values were used for machine learning analysis.

### Leaf Metabolomics

The metabolome represents a catalogue of small molecules present in a tissue, likely stemming from metabolic processes (Oliver *et al*., 1998; Tweeddale *et al*., 1998). Metabolomic analysis was completed at veraison and harvest on the 72-vine set. Three mature leaves were sampled from the middle of a shoot and immediately flash frozen in liquid nitrogen to capture the metabolic state of the leaves when attached to the vine. Frozen leaves were transported to the University of Missouri Enology lab on dry ice and stored at −80°C. Leaf metabolomes were analyzed using a modified form of a previously established protocol (Islam *et al*., 2011); Supplemental Note 2). LC-MS instrument files were converted to .cdf format and uploaded to XCMS online (Tautenhahn *et al*., 2012) for chromatogram normalization and feature detection via “single job” parameters. Identified metabolomic features were used as the basis of a principal components (PC) analysis. The top 20 PCs were treated as distinct phenotypes to model according to the experimental design. In PCs that varied significantly by rootstock, features that loaded more than 1.96 standard deviations above or below the mean were fit independently with the same model design.

### Gene Expression

The youngest fully-opened leaves (~1 cm) on two shoots were collected from each plant of the 72-vine set and pooled for RNA sequencing. Samples were sequenced using 3’-RNAseq, a method ideal for organisms with reasonably characterized reference genomes (Tandonnet & Torres, 2017). The first 12 nucleotides from each read were trimmed to remove low-quality sequences using Trimmomatic (options: HEADCROP: 12; (Bolger *et al*., 2014)). Low quality trimmed reads were additionally identified based on overrepresentation of kmers and removed using BBduk (April 2019 release) (Bushnell, 2017). Trimmed and QC-controlled reads were mapped to the 12Xv2 reference *Vitis vinifera* genome (Jaillon *et al*., 2007; Canaguier *et al*., 2017) using STAR (v2.7.2b) (Dobin *et al*., 2013) with default alignment parameters. RNAseq read alignments were quantified using HTSeq-count (v0.11.2) (Anders *et al*., 2010) and a modified version of the VCost.v3 reference *V. vinifera* genome annotation (Canaguier *et al*., 2017). To capture mis-annotated gene body boundaries in the genome, all gene boundaries in the annotation were extended 500 bp.

Variation in gene expression was assessed using two methodologies. First, we identified individual genes which responded to specific factors in the experimental design using DESeq2 (Love *et al*., 2014). Genes were filtered to a gene set that included only genes with a normalized count greater than or equal to two in at least five samples. Each filtered gene was fit with the model “~ Block + Irrigation + Phenology_Rootstock” where the ‘Phenology_Rootstock’ model term was used to understand the potential interaction of phenology and rootstock. Differentially expressed genes were identified for each pairwise contrast in the model. Second, we used principal component analysis (PCA) to identify co-expressed genes and analyzed the top PCs in the context of the broader experiment. Filtered genes were transformed using the variance stabilizing transformation (VST; (Anders & Huber, 2010)) and input into a PCA. To approximate the impacts of both spatial variation and pseudotime (row) in the vineyard, linear models were first fit to remove variation imparted by irrigation for each of the top 100 PCs. The residuals from these models were then used as the basis for linear models and as the basis for machine learning analysis.

### Leaf Shape

All leaves from a single shoot directly emerging from a trained cordon were collected from each vine at 80% anthesis and veraison. At harvest, we collected only the oldest (first emerging leaf), middle (estimated from the middle of a whole shoot), and youngest (smallest fully emerged leaf at the shoot tip, >1cm). Leaves were collected approximately in row order (from south to north) and stored in a cooler. Each leaf was imaged using an Epson DS-50000 scanner. In order to mimic the sampling regime at harvest, we subset the leaves collected at anthesis and veraison by extracting the youngest leaf, the approximate middle leaf, and the oldest leaf sampled.

We assessed leaf morphological variation using generalized procrustes analysis (GPA) of landmarks. For each leaf, 17 homologous landmark features were identified (Chitwood *et al*., 2014). The GPA-rotated coordinate space was used for all subsequent statistical analysis including PCA in order to summarize variation in leaf shape (Dryden & Mardia, 2016). From the PCA, we extracted the top 20 PCs and fit linear models and machine learning models to describe variation.

### Vine physiology

Intracellular CO_2_ concentration, stomatal conductance and leaf transpiration rate were measured on a fully expanded sun-exposed leaf during midday (10 am to 1 pm) using an LI-6400XT Portable Photosynthesis system coupled with a pulse amplitude-modulated (PAM) leaf chamber fluorometer with the following parameters: incident photosynthetic photo flux density level of 1000 μmol m-2 s-1 generated by a red LED array and 10% blue light to maximize stomatal opening (Li-Cor, Inc., Lincoln, NE, USA), CO_2_ mixer of 400 umol/s, fixed flow of 300 umol/s, and ambient leaf and block temperature. Soil moisture was measured for each plant in the 72-vine set using a fieldScout TDR 300 Moisture meter equipped with 20 cm rods (Spectrum Technologies, Inc. Aurora, IL, USA). Midday stem water potential was measured using a pressure bomb/chamber (PMS Instrument Co., Albany, OR, USA) after enclosing the leaves in an aluminum foil bag for at least 15 minutes to equilibrate the water potential of the xylem in the stem to that of attached leaf.

### Linear Models

Linear models were fit to the 20 measured ion concentrations, the top 20 PCs of the leaf metabolome, the top 100 PCs of the leaf transcriptome, the top 20 PCs of leaf morphospace, and each measured physiological trait. Each model was fit with fixed effect factors representing phenological stage (anthesis, veraison, or harvest), rootstock (Ungrafted, ‘1103P’, ‘3309C’, or ‘SO4’), leaf position (youngest, middle, or oldest; only used in leaf morphology and leaf ion concentration models), and all pairwise interactions of those terms. Both irrigation and block were included as fixed, non-interacting effects with the exceptions of physiology and metabolomics, for which we allowed the interaction of ‘Block’ as it correlates with the time of sampling. Row, an additional correlate for time and spatial variation, was included in place of a temporal block for the gene expression models after removal of the variation attributable to irrigation, a factor collinear with row. All linear models were interpreted using a type-3 sum of squares computation using the R package ‘car’ (Fox *et al*., 2013). Estimated p-values for each term in the models were corrected for multiple tests (within phenotype) using FDR correction as implemented by the R package ‘stats’ (R Core Team, 2013). Results from the models are reported as the variation explained by a particular term in the model and the estimated p-value. When appropriate, post-hoc mean comparisons were computed using the package ‘emmeans’ (Lenth *et al*., 2018). Where multiple linear models were being simultaneously interpreted, we applied a Bonferonni correction to reduce the number of false positives.

### Machine Learning to Identify Rootstock Effects

For visualization of between-class variation, we fit linear discriminant analysis models (LDA) to the full phenotypic data sets of ionomics, metabolomics, gene expression, and leaf morphology using the ‘lda’ function of the R package ‘MASS’ (Ripley, 2002). Projections of all samples into the LD space were plotted using ggplot2 (Wickham, 2016). In addition, we employed machine learning to capture subtle experimental effects. We partitioned phenotypic data sets into 80% training partitions and 20% testing partitions. Models were fit to predict the phenological stage from which a sample was taken, the rootstock to which the scion was grafted, and the joint prediction of phenology and rootstock. We also tested the predictability of leaf position for ionomics and leaf shape, and the interaction of rootstock and leaf position for ionomics. We used the ‘randomForest’ (Liaw *et al*., 2002) implementation of the random forest algorithm. Models were fit and tuned using the R package ‘caret’ (Kuhn, 2013). Each performance was assessed using accuracy, with performance on each class being assessed using the balanced accuracy, the midpoint of class-wise sensitivity and specificity. Where appropriate, models were compared to ‘chance’, or the occurrence frequency of each class. Confusion matrices were visualized from the out-of-bag predictions using ggplot2. Important features were identified from the randomForest object based on a phenotype-specific mean decrease in model accuracy (MDA).

### Phenomic trait covariation

We extracted from each data set the youngest available leaf from the 72 vine-set from which ionomics, metabolomics, gene expression, and leaf shape were measured. Each class of phenotypic data was summarized along the primary dimensions of variation using PCA. For each class, we extracted the top 10 PCs and fit Pearson’s correlations across all pairs of PCs at each phenological stage. P-values from computed correlations were corrected using the FDR method from the package ‘stats’ (Team & Others, 2013). Correlations and their strengths were visualized using the R package ‘igraph’ (Csardi *et al*., 2006). Example correlations were reported after running 10,000 bootstrapped subsamples of 90% of data for paired traits. From the distribution of estimated correlation coefficients, confidence intervals were computed from the 0.025 and 0.975 quantiles. A subset of example correlations were plotted using the R package ‘ggplot2’ (Wickham, 2016).

## Results

### Leaf ionome

To characterize the leaf ionome over the growing season, we sampled the youngest, middle, and oldest leaf on a single shoot from each of 288 vines at three phenological stages for ionomics analysis (Fig. 1). Bivariate correlations showed that ion concentrations are not independent of each other. However, the strength and direction of relationships between ions vary with respect to most experimental factors (for example, phenological stage and leaf position; Sup Fig. 1). As such, we fit independent linear models to each ion. Leaf position, phenological stage, or the interaction of phenological stage and leaf position explained the highest amount of variation for most ions (Fig. 1a-b). Many ions significant for the interaction showed a clear signal of leaf position at anthesis and veraison, and either no explainable variation or muted variation at harvest. For example, calcium (Fig 1b) varied with leaf position (22.7%; p < 1e-05), phenology (24.0%; p < 1e-05), and their interaction (7.4%, p < 1e-05). All possible pairwise combinations of leaf position were significantly different at anthesis, and both the youngest and middle leaves were different from the oldest leaves at veraison and harvest. In the case of potassium (Fig 1b), significant variation was explained by leaf position (16.1%; p < 1e-05), phenology (19.6%; p < 1e-05), and their interaction (10.6%; p < 1e-05). However, post-hoc comparisons showed that differences were present only at anthesis and veraison. Ions that responded weakly to the interaction of leaf position and phenology tended to show significant variation explained by the interaction of rootstock and phenology (see below). These ions showed similar patterns to the leaf position by phenology interaction where clear signal is exhibited at anthesis and veraison then is either absent or muted at harvest (see, for example, cobalt and nickel; Fig 1c).

**Figure 1.**
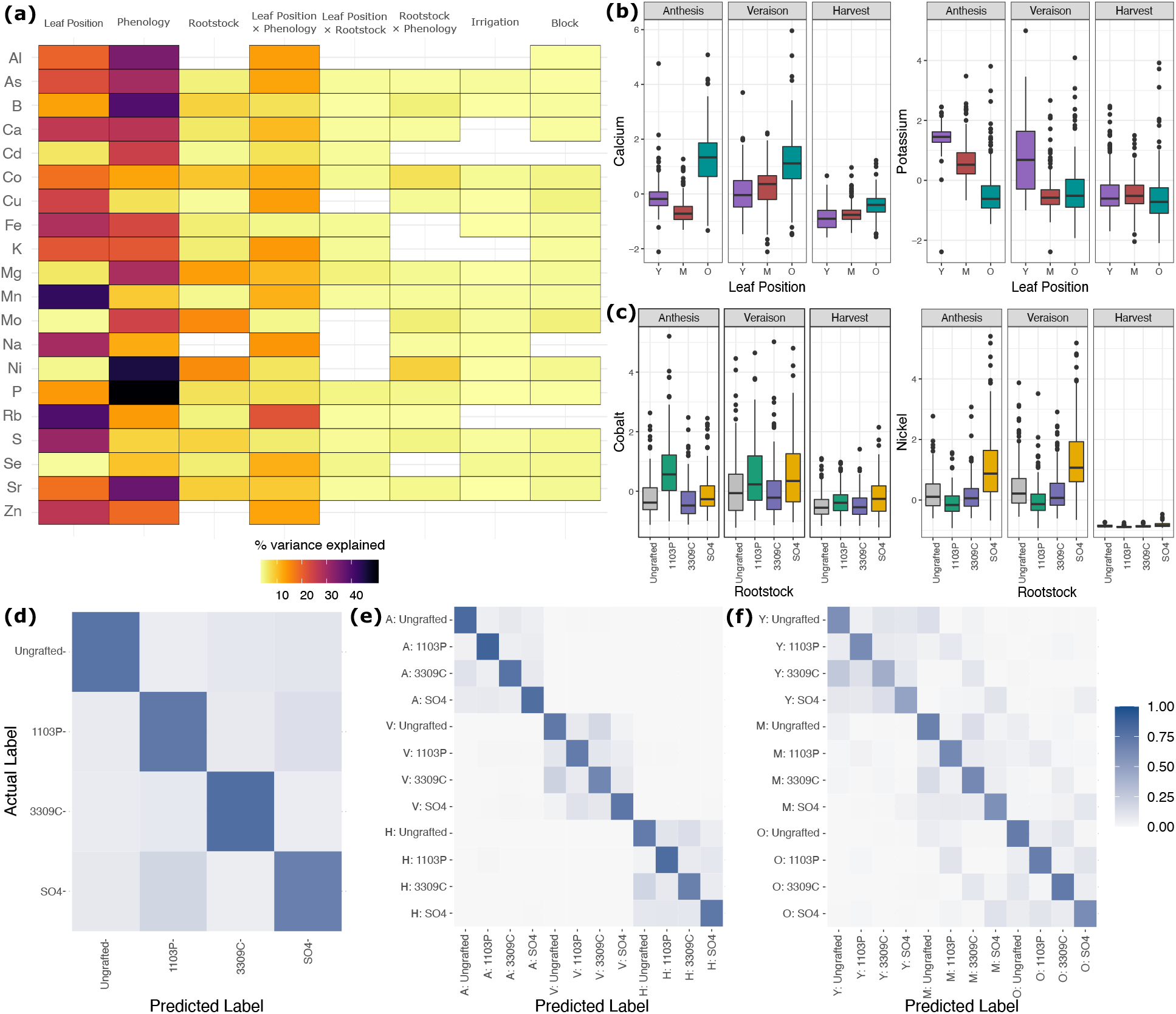
The ionome shows strong signal from rootstock genotype, leaf position, and phenological stage. **(a)** Percent variation captured in linear models fit to each of 20 ions measured in the ionomics pipeline. Presence of a cell indicates the model term (top) was significant (FDR; p.adj < 0.05) for that ion (left). **(b)** Example ions shown to vary significantly by the interaction of leaf position and phenological stage. Boxes are bound by 25th and 75th percentile with whiskers extending 1.5 IQR from the box. (**c**) Example ions shown to vary significantly by the interaction of rootstock genotype and phenological state. Boxes are bound by 25th and 75th percentile with whiskers extending 1.5 IQR from the box. (**d**) Standardized heatmap for out-of-bag (OOB) predictions by a random forest trained to predict rootstock genotype, **(e)** the interaction between rootstock genotype by phenology, and **(f)** the interaction between rootstock genotype and leaf position.

Machine learning on ion concentrations showed that rootstock and the interactions of rootstock with phenology and leaf position were independently predictable classifications. A random forest model trained to predict rootstock showed an overall accuracy of 75.2% (Fig 1d). Ions important for this classification were nickel (MDA=0.089), molybdenum (MDA=0.058), and magnesium (MDA=0.054). Notably, when we trained a model to simultaneously predict phenological stage and rootstock, rootstock prediction accuracy increased appreciably (Fig. 1e). For example, the ability of the model to detect ungrafted vines (the balanced accuracy of ungrafted predictions) improved from 81.7% accuracy overall to 91.1% accuracy at anthesis and 85.9% at harvest. Generally, performance at veraison matched the rootstock-only model performance. The ions most important for this simultaneous classification were nickel (MDA=0.167), phosphorus (MDA=0.110), and strontium (MDA=0.065). Interestingly, the joint prediction of rootstock and leaf prediction performed substantially better than chance (p < 1e-05), however average performance of the model as assessed through class-wise balanced accuracies were comparable to if not slightly worse than just predicting rootstock (Fig 1f). Ions important for this classification were sulfur (MDA = 0.051), rubidium (MDA = 0.051), and nickel (MDA = 0.049).

### Leaf metabolomics

We performed untargeted metabolomics on leaves from 72 vines at veraison and harvest, quantifying the concentrations of 661 metabolites (Fig. 2). The top 20 PCs accounted for a total of 67.3% of the total metabolomic variation, with the top three capturing 23.1%, 9.2%, and 6.2%, respectively. Linear models for each of the top 20 PCs found that the strongest drivers of variation in leaf metabolomics were phenology and temporal blocking factor. For example, 90.6% of variation on PC1 was due to phenology (p < 1e-05; Fig 2a). PC2 primarily reflected the interaction of phenology and temporal block (26.4%, p < 1e-05) and temporal block as a main effect (18.9%, p < 1e-05). The patterns of variation attributable to PC2 were similar in PCs 3-10 (Fig 2a).

**Figure 2.**
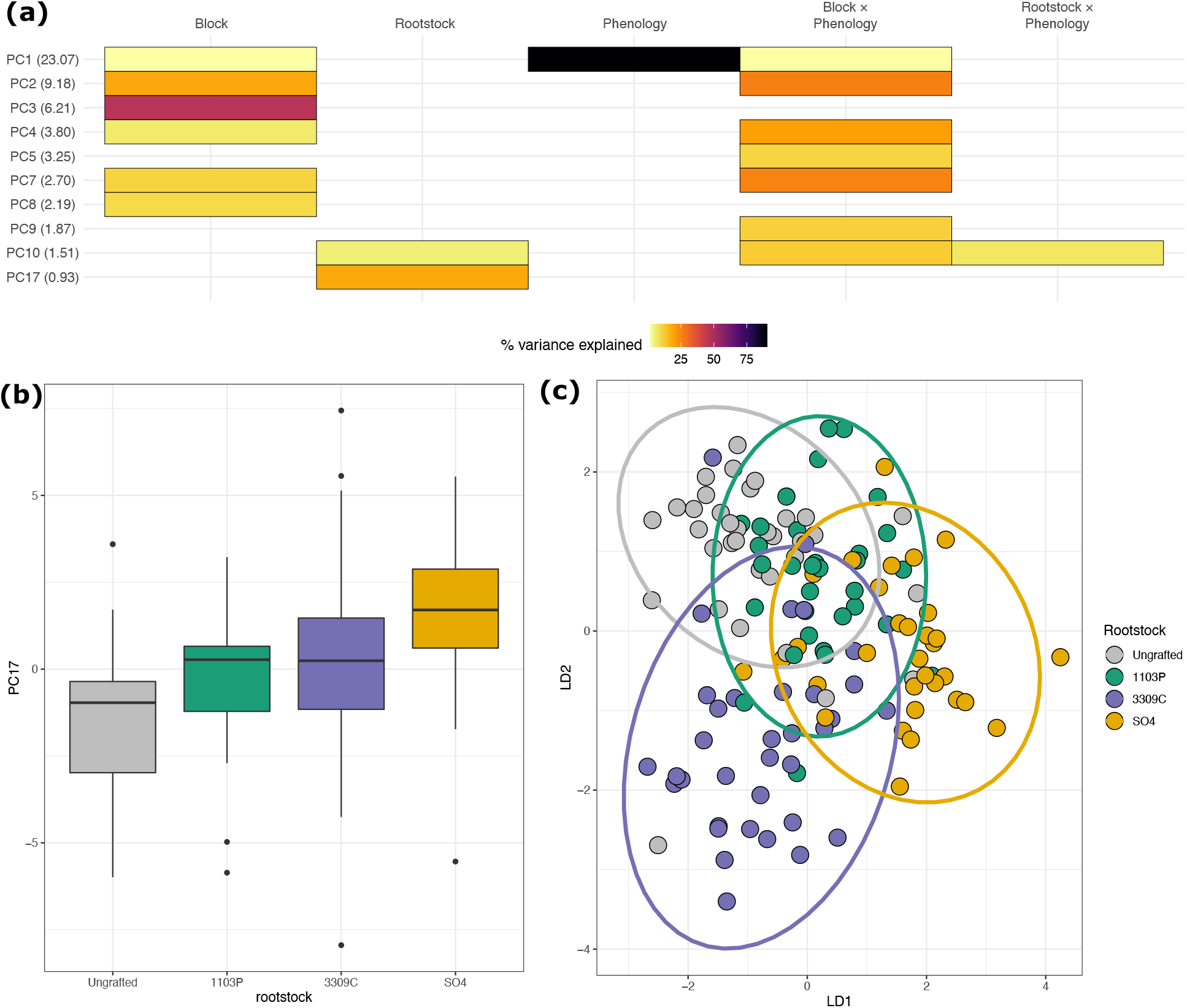
The metabolome is influenced by rootstock genotype, phenological stage, and time of sampling. **(a)** Percent variation captured in linear models fit to each of the top 20 principal components of the metabolome (661 measured metabolites). Presence of a cell indicates the model term (top) was significant for that PC (left, percent variation explained by the PC in parentheses). **(b)** The distribution of projections onto PC17, the strongest captured rootstock effect in the metabolome. Boxes are bound by the 25th and 75th percentiles with whiskers extending 1.5 IQR from the box. **(c)** Projections of all samples into the first two dimensions of a linear discriminant space trained to maximize variation between rootstock genotypes.

PC17 was controlled by rootstock as a main effect (18.5%, p < 1e-03; Fig 2b). On PC17, ungrafted vines were significantly different from vines grafted to ‘3309C’ (p = 0.02) and ‘SO4’ (p < 1e-05). Vines grafted to ‘ 1103P’ were also significantly different from vines grafted to ‘SO4’ (p = 0.009). Metabolites that loaded more than 1.96 sd from the mean loading on PC17 were extracted and independently fit to additional linear models. We identified four metabolite features (M374T1 [rt = 1.33, m/z = 374.1146], M117T1 [rt = 0.61, m/z = 117.0583], M175T1_1 [rt = 0.87, m/z = 175.1269], and M333T1_3 [rt = 0.71; m/z = 333.1582]) which were influenced by rootstock as a main effect and the metabolite (M112T1 [rt = 1.48, m/z = 112.0061]) which was influenced by the interaction of rootstock genotype and phenological stage.

Linear discriminant analysis confirmed that many experimental factors likely influence the metabolome. For example, when trained to maximize variation between classes of rootstocks, the model identified a space that weakly separates ‘1103P’-grafted and ‘SO4’-grafted vines from Ungrafted and 3309C-grafted vines (LD1) and separates 3309C-grafted vines from other classes (on LD2) (Fig 2c). Despite this, machine learning showed minimal predictability for any class other than phenology, which was predictable with an accuracy of 100% for withheld samples. Rootstock genotype was not predictable with accuracy only marginally better than chance (34.6%).

### Gene Expression

We performed 3’-RNAseq on 72 vines at three time points (Fig. 3). We identified variation in 23,460 genes that had a DESeq2-normalized count greater than two in at least five samples. Hierarchical clustering of the 500 most variable genes after variance stabilizing transformation (VST) showed that most variation in the transcriptome was explained by phenological stage (Fig 3a). The top 100 PCs on the VST-transformed gene counts accounted for nearly 92.3% of variation in the transcriptome. Linear models on each of the top 100 PCs indicated that 82.4% and 61.4% of the variation on PC1 and PC2 respectively were attributable to the phenological stage (Fig 3b-c). Row was also a significant descriptor of variation as a single, fixed effect and in interactions with rootstock and phenological stage. For example, row accounted for 36.0% and 43.3% of the variation on PC4 and PC6, respectively. Interacting with phenological stage, row accounted for >10% of variation on 17 additional PCs.

**Figure 3.**
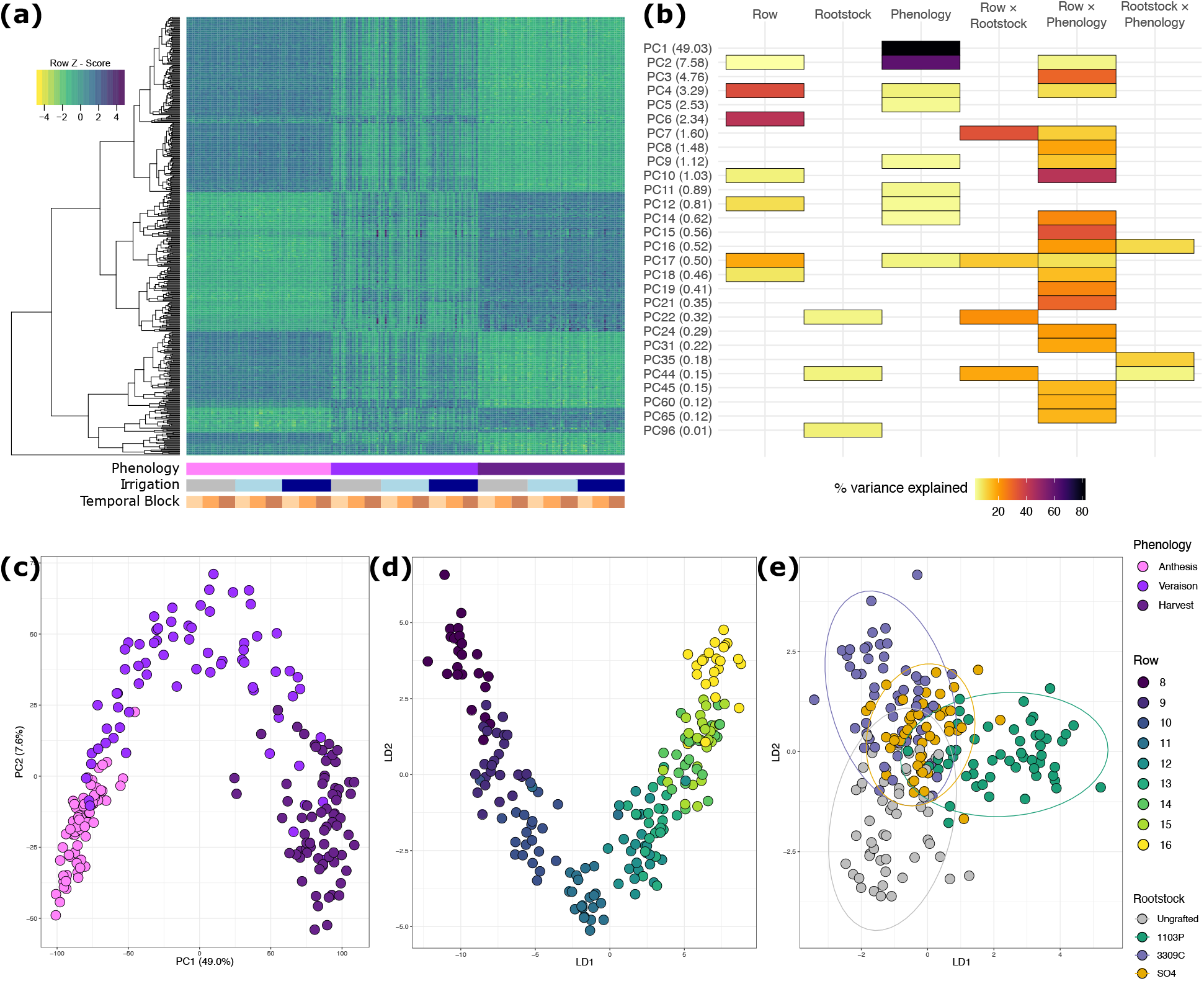
Gene expression primarily responds to time of season and circadian correlates. **(a)** Heatmap showing 500 genes with the highest variance following the filtering of lowly expressed genes and gene-by-gene variance stabilizing transformations (VST) ordered by example model factors (below). **(b)** Percent variation captured in linear models fit to the top 100 Principal Components of the VST-transformed gene-expression space. Presence of a cell indicates the model term (top) was significant for that PC (left, percent variation explained by the PC in parentheses). **(c)** Projections of all samples projected into the first two dimensions of the linear discriminant space trained to maximize variation between phenological stages, **(d)** row of the vineyard, and **(e)** rootstock genotype.

LDA to separate phenological stages defined three distinct, non-overlapping groups in the space spanning LD1 and LD2 (Sup Fig. 2). When trying to separate rows into distinct classes, the model converged on a ‘horseshoe’ shape in the LD1-LD2 space (Fig 3d). LD1 maximized the variation between row 8 (sampled early in the day) and row 16 (sampled a few hours later). LD2 maximized the separation of both rows 8 and 16 with row 12 (the row sampled in the middle of the sampling window). A model trained to separate rootstock classes (Fig. 3e) showed that LD1 separated the rootstock 1103P from other rootstock genotypes, and LD2 primarily separated the rootstock ‘3309C’ from ungrafted vines (Sup Fig. 2).

Formal machine learning on gene expression PCs largely supported the linear models. A random forest trained to predict phenological stage classified testing samples with 92.9% accuracy. Anthesis was the most predictable class with a balanced accuracy of 100%; veraison and harvest displayed balanced accuracies of 92.7% and 92.4%, respectively. The PCs most important in phenology prediction were PC1 (MDA = 0.16) and PC2 (MDA = 0.12). Gene expression PCs were unable to predict rootstock, with a total prediction accuracy of 23.4%. While no features were especially important in the prediction processes, PC44 showed the largest mean decrease in Gini impurity corroborating its signal in the linear models.

### Leaf shape

We collected leaves from the 288-vine set at three time points and landmarked a total of 2,422 leaves (Fig. 4). Homologous leaf landmarks were used for generalized procrustes analysis (GPA). PCA on the GPA-rotated coordinates revealed ~97.2% of the total shape variation was captured by the top 20 principal components with PC1, PC2, and PC3 explaining 24.1%, 19.0%, and 13.3% of the variation respectively. Lower values on PC1 primarily capture leaves with shallow petiolar sinuses and short midvein distance from the depth of the superior sinus to the top of the midvein, whereas higher values on PC1 capture the opposite (Fig. 4a). Similarly, lower values on PC2 capture deep petiolar sinuses combined with very shallow superior sinuses, and vice versa for higher values. PC3 primarily captures asymmetry (Fig. 4a).

**Figure 4.**
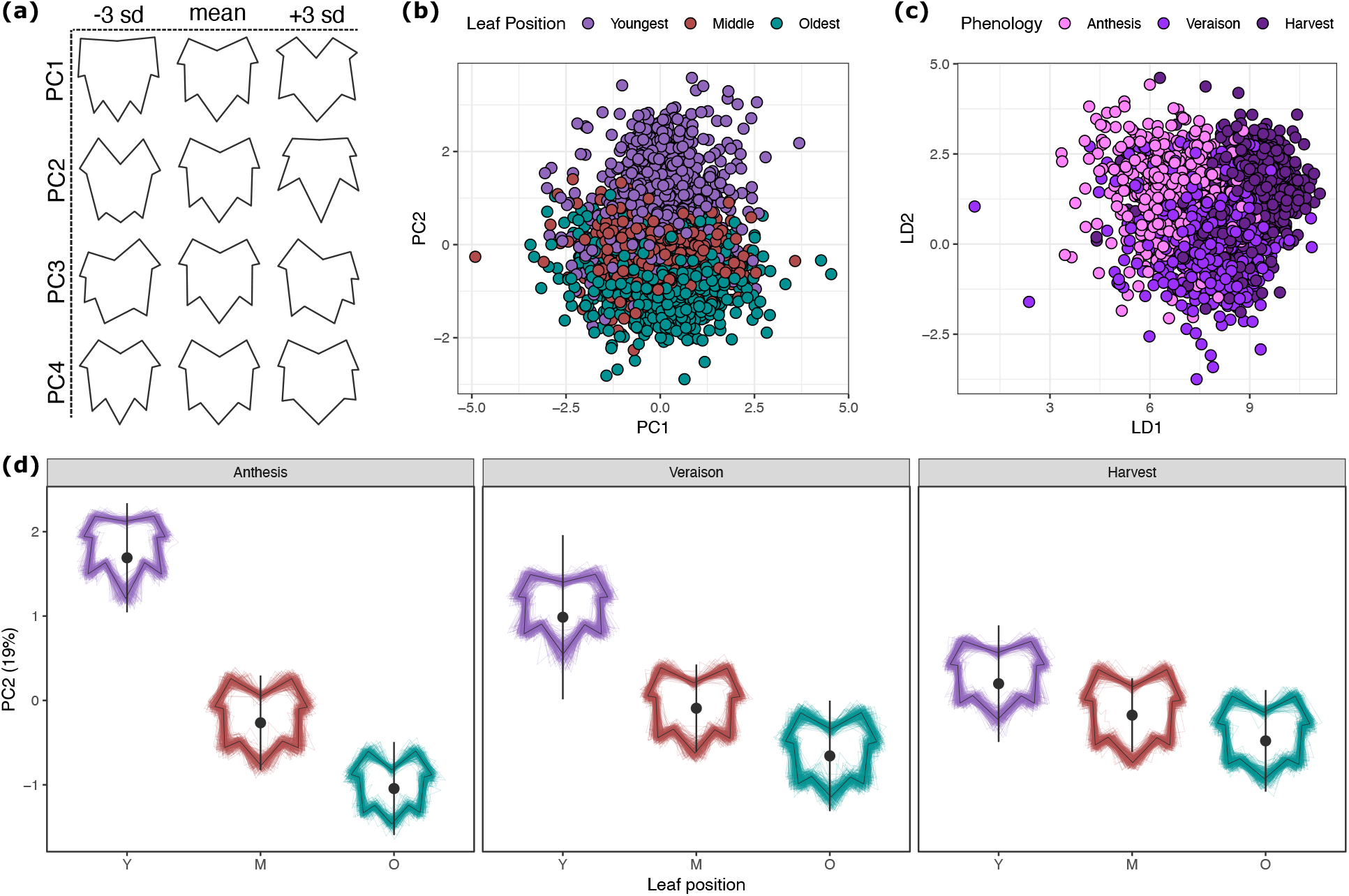
Leaf shape variation is primarily determined by shoot position but changes over the season. **(a)** Representative shapes showing leaf variation (−3 sd, mean, +3 sd) captured in each of the top 4 principal components of the Generalized Procrustes Analysis-rotated leaf shapes. **(b)** Projections of all leaves into the first two dimensions of principal component space colored by the strongest determinant of variation in the top two PCs. **(c)** Projections of all leaves into the first two dimensions of a linear discriminant space trained to maximize variation between phenological stages. **(d)** Variation in leaf shape captured on PC2 shown by leaf position and phenological stage. Large points represent the mean of the group when projected onto PC2. Bars surrounding the mean show one standard deviation. Variation in each group is shown as a composite leaf trace scaled to a standard size and centered over the mean.

In total, only 5.76% of variation on PC1 was explained by the experimental design, with most variation explained by phenology (2.63%; padj < 1e-05), rootstock (0.95%; padj < 0.001), leaf position (2.61%; padj = 0.03), and the interaction of phenology and leaf position (0.62%; padj = 0.009) (Sup Fig 3a). Post-hoc mean comparisons on PC1 showed that shapes of leaves from ungrafted vines were significantly different from leaves of vines grafted to 3309C (p < 0.001) and SO4 (p < 0.001) (Sup Fig 3b). Moreover, PC1 captured subtle variation in the leaf position by phenological stage interaction where middle leaves showed significant differences between anthesis and veraison (p < 1e-03), and the oldest leaves showed significant differences when comparing anthesis to veraison (p < 1e-05) and anthesis to harvest (p < 1e-03).

For PC2, 61.4% of variation could be assigned to an experimental factor. This included significant variation from leaf position (46.9%, padj < 1e-05), phenology (1.4%; padj < 1e-05), and the interaction of leaf position and phenology (12.05%; padj < 1e-05; Fig 4d). Specifically, younger leaves tended to have shallower sinuses and exaggerated superior sinus depths (higher values on PC2), whereas older leaves tended to develop deeper petiolar sinuses and more shallow superior sinuses (lower values on PC2). The degree of this separation decreased across the season, and the shapes converged on the mean leaf shape on PC2, consistent with the middle leaf at all three phenological stages. PC2 additionally reflected the interaction of leaf position and rootstock (0.22%; p = 0.04; (Sup Fig. 4b)), but post-hoc comparisons did not find any significant pairwise comparisons.

Machine learning on the GPA-rotated coordinate space identified moderate division of developmental and phenological classes. Random forest models could predict the leaf position with 73.1% accuracy, with the most important feature being the y-component of the leaf apex (MDA = 0.051). A model trained to predict phenology performed at 64.3% with the most important features being the x-components of the points corresponding to superior sinus depth (left sinus MDA = 0.030, right sinus MDA = 0.019). A model trained to predict rootstock performed only marginally better than chance at 28.1% accuracy.

### Vine physiology

For 72 plants in the vineyard, we measured intracellular CO_2_ concentration (C_i_), stomatal conductance (gs), leaf transpiration, water potential (*ψ*), and soil moisture (Fig. 5). Each physiological trait varied significantly across phenology and the block by phenology interaction (Fig 5a). For example, at harvest, we observed specific differences in leaf CO_2_ concentration (A vs C: p=0.003; B vs C: p=0.002) and leaf transpiration (A vs B: p < 1e-03; A vs C: p < 1e-05; B vs C: p < 1e-05). Additionally, stomatal conductance and leaf transpiration rate varied significantly with the interaction of rootstock and phenology. For both traits, a post-hoc comparison of means showed that these values were elevated in 1103P at veraison as compared to ungrafted vines (stomatal conductance: p = 0.002; leaf transpiration: p = 0.001; Fig 5b-c).

**Figure 5.**
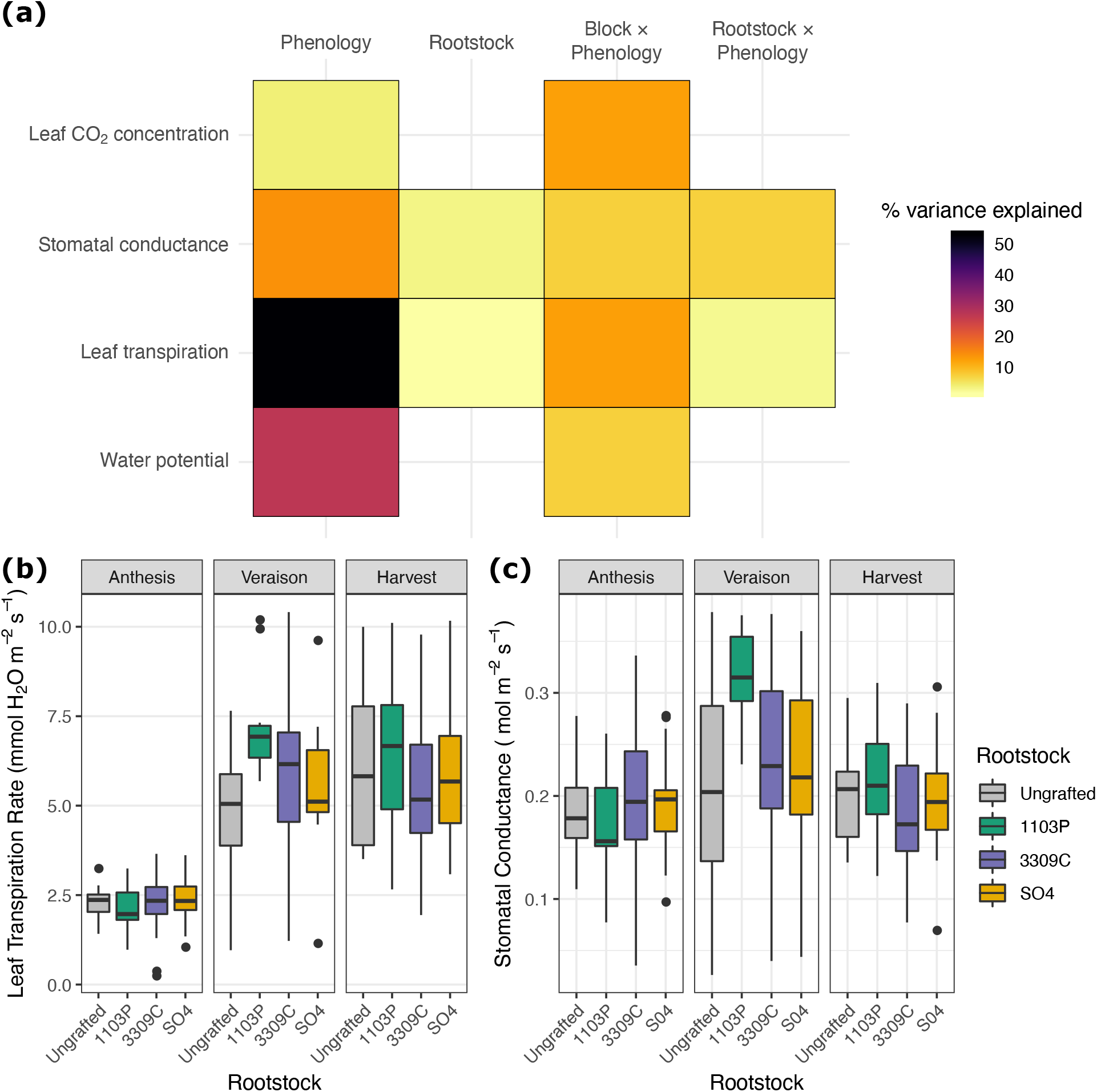
Vine physiology measurements show signal from most experimental manipulation. **(a)** Percent variation explained by model terms (top) from linear models fit to each of four physiology traits (left). **(b)** Variation in leaf transpiration rate for each rootstock genotype over the course of the season. Boxes are bound by the 25th and 75th percentiles with whiskers extending 1.5 IQR from the box. **(c)** Variation in stomatal conductance for each rootstock genotype over the course of the season. Boxes are bound by the 25th and 75th percentiles with whiskers extending 1.5 IQR from the box.

### Phenomic trait covariation

For each of the 72 plants measured for all phenotypes in the vineyard, we explored the extent to which different phenotypes covaried (Fig. 6). Within each phenotyping modality, we summarized the primary dimensions of variation using PCA. From each PCA, we extracted the top ten PCs, which explained a total of 88.9% of variation in the ionomics PCA (iPCA), 55.9% of the variation for the metabolomics PCA (mPCA), 74.8% of the variation in the gene expression PCA (gPCA) and 87.9% of the variation in the leaf shape PCA (sPCA). Pairwise correlations of each PC within each phenological stage showed diverse correlation magnitudes and directions both within a phenotyping modality and between phenotyping modalities (Fig 6a-c). Generally, the strongest relationships were between PCs within phenotypic modalities. For example, the strongest correlations identified were between gPC1 and gPC2 at anthesis (r = 0.85, CI = [0.81, 0.87]; Sup Fig 4a), and mPC1 and mPC2 at harvest (r = −0.78, CI = [−0.82. −0.76]). Correlations between modalities represented a diversity of responses across phenological stages. For example, the correlation between gPC4 and sPC3 is similar across the phenological stages, but only the correlation at veraison is significant (r = 0.41, CI = [0.34, 0.47]; Sup. Fig 4b). Correlations such as between mPC3 and gPC6 were similar and significant at both veraison (r = −0.44, CI = [−0.50, −0.37]; Sup Fig 4c) and harvest (r = −0.37, CI = [−0.45, −0.28]; Fig 6c). While many correlations varied over the course of the season, some relationships entirely shifted in direction. For example, the correlation between mPC3 and mPC6 shifted from a positive significant relationship (r = 0.58, CI = [0.52, 0.63]) at veraison to a negative significant relationship at veraison (r = −0.66, CI = [−0.73, −0.59]) (Sup Fig 4d).

**Figure 6.**
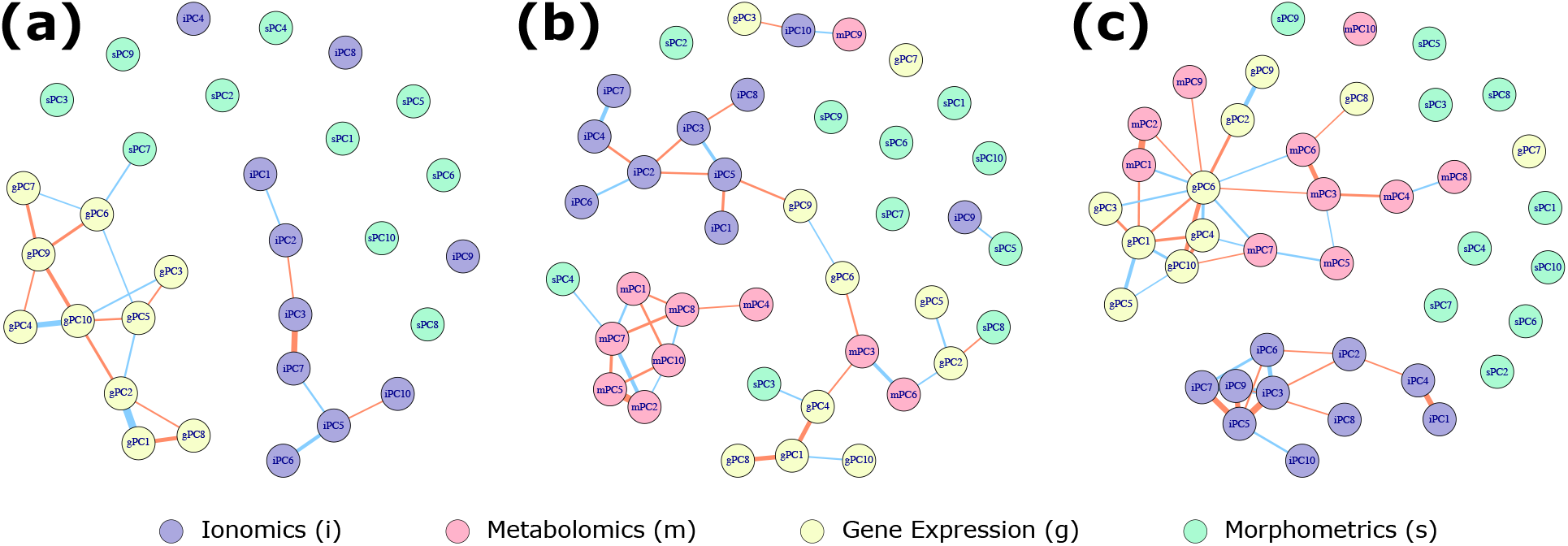
Trait covariation varies over the course of the season. Correlation networks showing patterns of covariation within and between phenotyping modalities. Nodes of the network are connected if they are significantly correlated (Pearson, FDR; p.adj < 0.05). Edge thickness is proportional to the strength of correlation (multiplied by 16 for visibility). Edge color reflects the direction of the correlation where blue edges indicate positive correlations and orange edges indicate negative correlations. Modalities are indicated by a leading character and node color: ionomics (iPCs; purple), metabolomics (mPCs; pink), gene expression (gPCs; yellow), leaf shape (sPCs; green). Network topologies are shown for **(a)** anthesis, **(b)** veraison, and **(c)** harvest.

## Discussion

In this study, we characterized variation in leaf ionomics, untargeted metabolomics, transcriptomics, leaf morphology, and physiology in an experimental rootstock vineyard at three distinct time points over the course of a growing season. Overall, we find that time of season was the primary driver of most leaf phenotypic variation, and that rootstock influences on leaf traits can be season-specific. Generally, ‘Chambourcin’ leaves show subtle responses to grafting, with the strongest signals observed in phenotypes for which the root systems have a noted and well-understood role (e.g., ion concentrations in leaves).

### Phenology explains significant variation in all leaf phenotypes

We found that the phenological stage was the strongest driver of phenotypic variation for most leaf phenotypes. For example, all 20 ions varied with the phenology and most ions showed that phenology, or the interaction of phenology with leaf developmental position, was the strongest source of variation (Fig. 1). Additionally, nearly one third of all measured transcripts responded to seasonal variation, and the strongest effects on the transcriptome were the phenology and the row, a correlate for the time within a three hour sampling window. The only phenotype for which phenology was not the most explanatory factor is leaf shape. Consistent with previous studies (Chitwood *et al*., 2015), we confirm that most of the leaf shape variation measured reflects development along a single shoot, but much of this variation is explained via interaction with the phenology.

The seasonal component to grapevine phenotypic variation is a subject of much research, especially in the berry. In studies designed to characterize effects of cultivar variation and molecular underpinnings of terroir, seasonal variation was the strongest signal in the metabolome (Degu *et al*., 2014; Anesi *et al*., 2015; Cuadros-Inostroza *et al*., 2016; Dal Santo *et al*., 2016). Several studies have also sought to characterize transcriptomic variation over the course of the season. For example, in conjunction with metabolomics, seasonal variation of berry development was used to identify developmental markers in ‘Corvina’ (Zamboni *et al*., 2010). Follow-up analysis showed that nearly 18% of transcripts varied seasonally (Dal Santo *et al*., 2013). Grapevine leaves also vary tremendously in shape over the growing season (Chitwood *et al*., 2015) and are stable over multiple growing seasons; interestingly, the climate of the season in which the leaves were patterned influence aspects of leaf shape (Chitwood *et al*., 2016, 2020). We confirm that the patterns of variation previously identified in berries are also present in the leaves, and that patterns of leaf shape seem to be stable across studies.

While many studies have uncovered temporal effects on the ionome across years (Baxter *et al*., 2013; Pauli *et al*., 2018), variation within a single year or a single growing season remains relatively unstudied. One example included the joint analysis of the ionome and metabolome in Aleppo pine (*Pinus halepensis*), a perennial system with a bimodal growth habit in both spring and summer, where a suite of ions more abundant during spring growth were identified while only potassium was more abundant in the summer (López-Orenes *et al*., 2018). Other studies profiled tangential effects of the seasonal ionome; for example, winter-phased cultivars of barley (*Hordeum vulgare*) show differential uptake of nutrients in comparison to summer-phased cultivars, but the study was primarily targeted to identify genotypic rather than temporal effects (Thomas *et al*., 2016). Our data advances these previous studies by identifying the dynamic nature of ion uptake over the course of a season. More work is needed to understand how seasonal variation in ion concentrations vary inter-annually, by plant organ, or spatially; similarly, relationships between ion concentrations in leaves (a proxy for ion uptake) and berry chemistry and wine quality is another important area of future work.

### Grafting and rootstock genotype exhibits a complex and subtle signal on most leaf phenotypes

Consistent with previous studies, we confirm that grafting in general, as well as rootstock genotype, has a complex effect on phenotypic variation in grapevine shoot systems. Most notably, we show that the rootstock to which a scion is grafted is predictable from ion concentrations in the leaves, and that this signal is strengthened by inclusion of phenological stage. For example, we previously showed that nickel concentration was elevated in the rootstock ‘SO4’ (Migicovsky *et al*., 2019b). At a similar point in the season, we observe the same pattern, but by harvest, nickel is almost entirely excluded from the leaf suggesting that the biological implications of this differential uptake could be missed if not surveyed across the season. We also confirm that rootstock genotype influences the metabolome of grafted grapevine, in some cases in a season-specific manner. In the transcriptome, PCA was able to identify dimensions of variation that were significantly described by rootstock and the interaction of rootstock and time of day, confirming prior observations (Migicovsky *et al*., 2019a). Moreover, supervised methodologies identified linear discriminants in the PC space that weakly separated some rootstock genotypes. However, gene-by-gene analysis (with default p-value correction regimes) finds no genes modulated by rootstock genotype, or even just from the act of grafting. Finally, of the physiology traits we measured, leaf transpiration and stomatal conductance were higher in ‘1103P’ in the middle of the season. Thus the impact of grafting on leaf phenotypic variation varies by phenotype. Regardless, we identify subtle but ubiquitous effects from rootstock genotype on shoot system phenotypes that are often season-specific.

The impact of root genotype on shoot phenotype is a growing area of research, especially in grapevine. For ‘Cabernet Sauvignon’, grafting increased ion uptake globally and some rootstock genotypes provide a clear signal in the scion (Lecourt *et al*., 2015; Gautier *et al*., 2020b). Also, the metabolome is a key driver of the formation of the graft junction and some key metabolites could be responsible for graft incompatibility (Canas *et al*., 2015). Building on this work, targeted metabolomics showed two classes of metabolites, flavanols and stilbenes, were differentially abundant at graft junctions and in the rootstocks of ‘Cabernet Sauvignon’ vines one month after grafting (Prodhomme *et al*., 2019). However, flavanols were not differentially abundant in the scion, but scion stilbene concentrations were apparently controlled by rootstock genotype. The effect of rootstock genotype on the scion transcriptome is perhaps the most varied. For example, ‘Cabernet Sauvignon’ shoot apical meristems show no effects by rootstock genotype (Cookson & Ollat, 2013), but berries of the same cultivar do, although the effect is tempered by seasonal variation (Corso *et al*., 2016). Variation in ‘Chambourcin’ leaf shape is also driven by rootstock genotype, especially in conjunction with differences in irrigation (Migicovsky *et al*., 2019a). Collectively, these studies all suggest that rootstock genotype influences scion phenotypes, but those effects will vary by phenotype, scion genotype, and perhaps other experimental conditions. Our results confirm this suggestion adding that aspects of time are tremendously influential to the observed results regardless of phenotype.

### Phenomic covariation warrants work toward latent phenotypes

In the present study, we assess the extent of covariation among leaf phenotypes. For the primary dimensions of variation in each data type, within-data-type correlations are strong. Correlations also exist between phenotypes, suggesting room for the analysis of latent phenotypic structure for experimental questions. For example, aspects of the metabolome are frequently correlated with other data types such as the transcriptome and aspects of leaf shape. Interestingly, correlations within and between data types are highly dynamic over a growing season. For example, several correlations with leaf shape were present at veraison, but were completely missing from anthesis and harvest. We believe this work warrants further investigation, specifically, by adding data on other phenotypic classes such as lncRNAs (Vitulo *et al*., 2014; Harris *et al*., 2017), epigenetics (Williams *et al*., 2020), and microbiomes (Marasco *et al*., 2018; Swift *et al*., 2020). Much of the work constituting phenomics in grapevine has addressed how berries develop over the growing season, how cultivars differ from one another, and how the concept of terroir influences wine (Zamboni *et al*., 2010; Palumbo *et al*., 2014; Degu *et al*., 2014; Anesi *et al*., 2015; Savoi *et al*., 2016, 2017). Despite data integration becoming more popular, there are still many open questions as to what methods are most appropriate and how to most effectively utilize them (reviewed for grapevine in (Wong & Matus, 2017; Fabres *et al*., 2017); reviewed broadly in (Huang *et al*., 2017; Stein-O’Brien *et al*., 2018). Ongoing work attempts to integrate high-dimensional phenotypic datasets generated within a single organ system (e.g., leaves); and future studies should expand this to explore phenomic variation in and among organs, over time, and across space.

## Supplemental Figures

**SFig 1.**
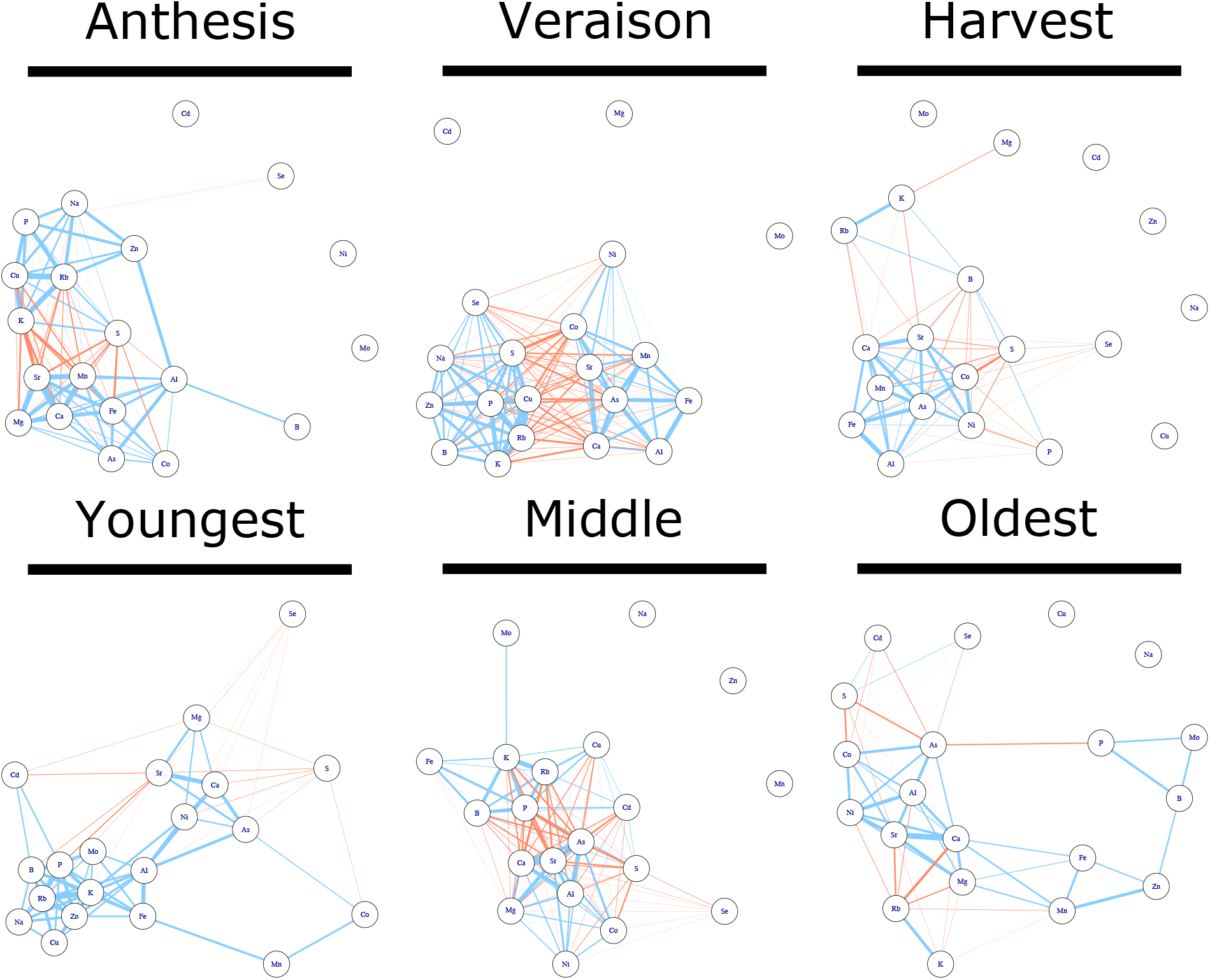
Patterns of ion covariation change over experimental treatments. Correlation networks showing patterns of ion covariation across phenological stages and shoot position. Nodes of the network are connected if they are significantly correlated (Pearson, FDR; p.adj < 0.05). Edge thickness is proportional to the strength of correlation (multiplied by 16 for visibility). Edge color reflects the direction of the correlation where blue edges indicate positive correlations and orange edges indicate negative correlations.

**SFig 2.**
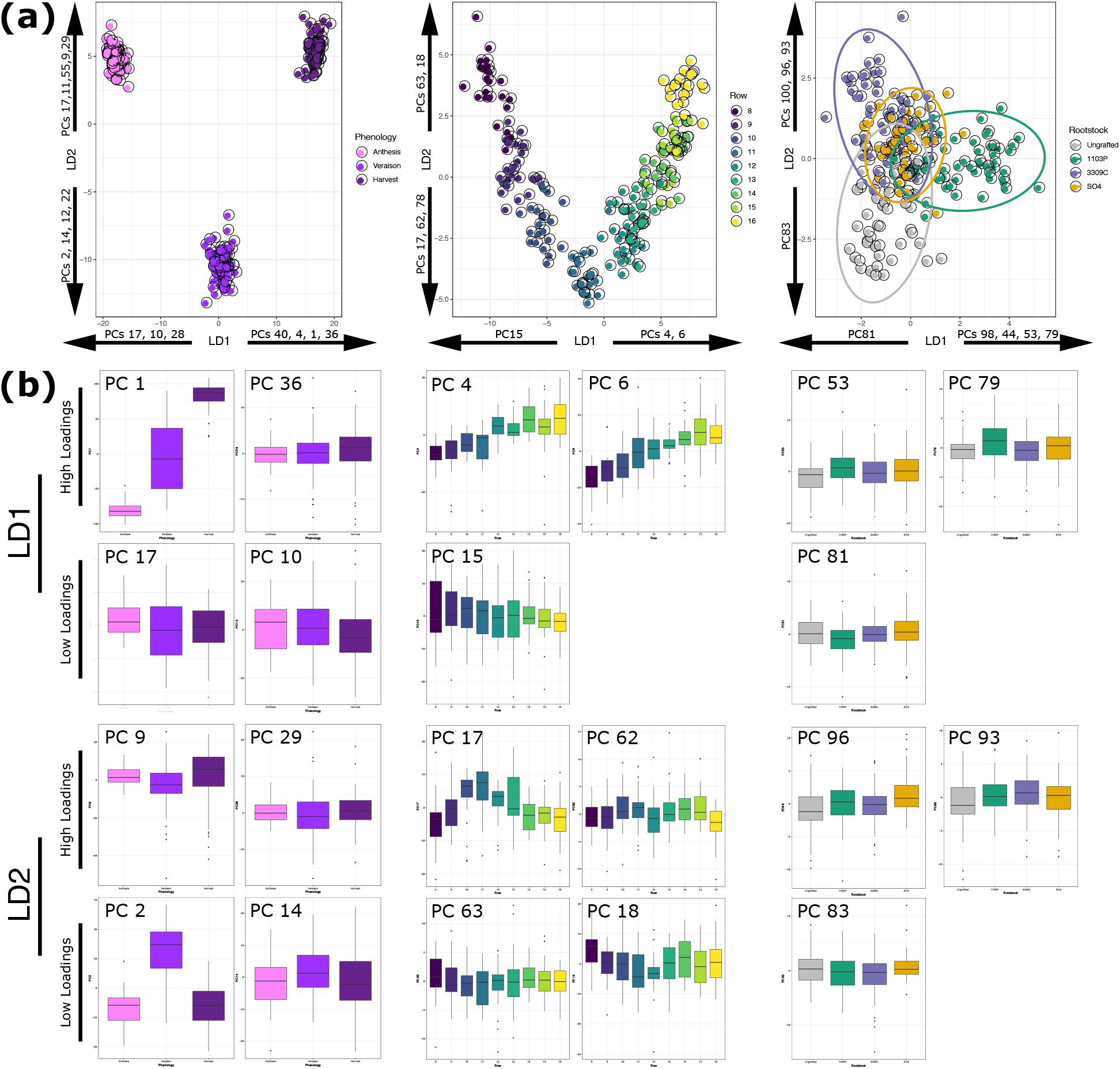
Patterns of variation contributing to gene expression linear discriminants. (A) Projections of leaf gene expression samples into the first two dimensions of a linear discriminant space trained to maximize variation between phenological stages, rows in the vineyard, and rootstock genotype. For each LD, the PCs that loaded significantly (>1.96 sd from the mean loading) are listed in order of loading magnitude. (B) Distribution of the top loading PCs onto LD1 and LD2 for each of the trained models.

**SFig 3.**
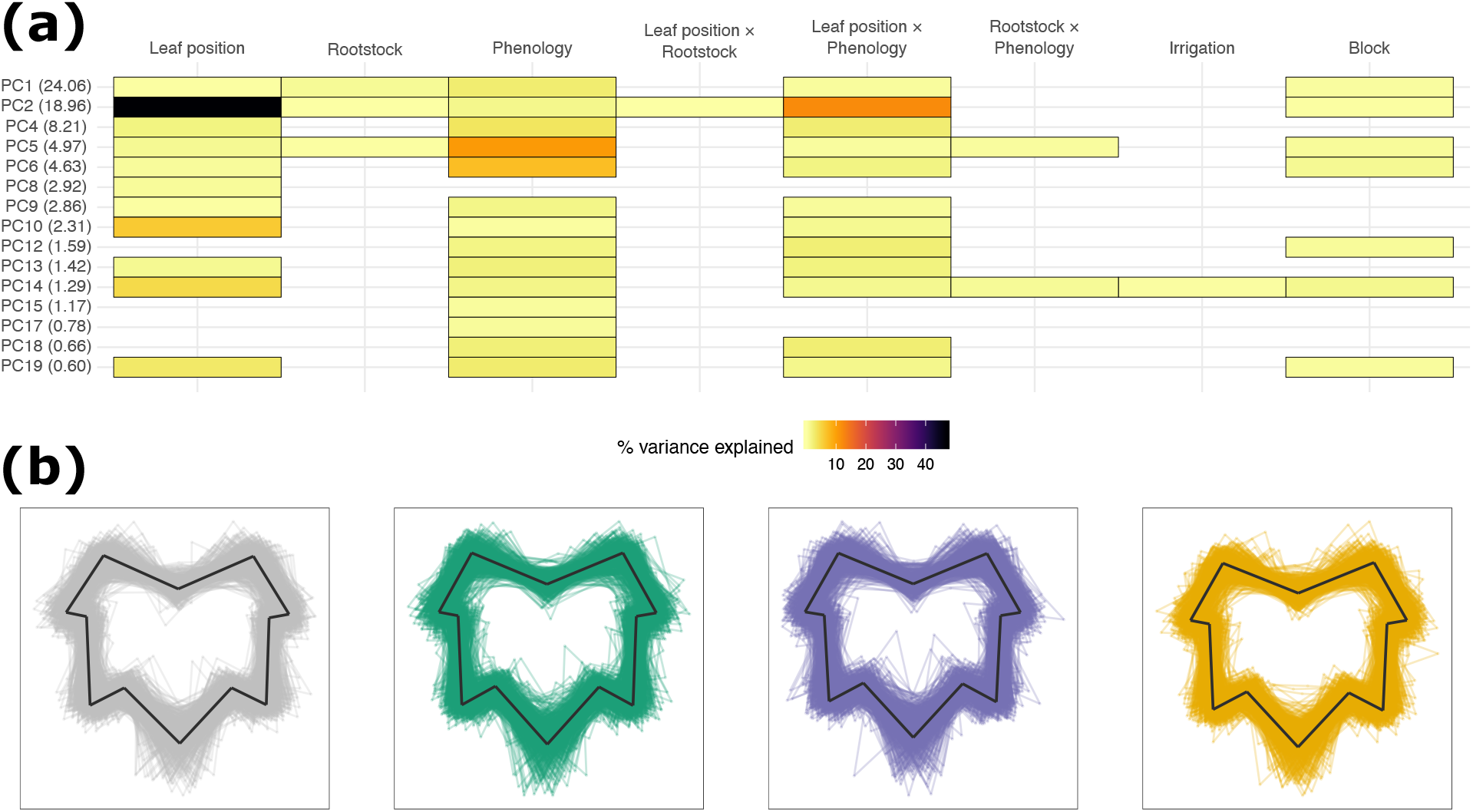
Patterns of variation in leaf are subtle. A Percent variation captured in linear models fit to each of the top 20 principal components of leaf morphology. Presence of a cell indicates the model term (top) was significant for that PC (left, percent variation explained by the PC in parentheses). (B) Composite leaf traces for the main rootstock genotype effect identified on PC1.

**SFig 4.**
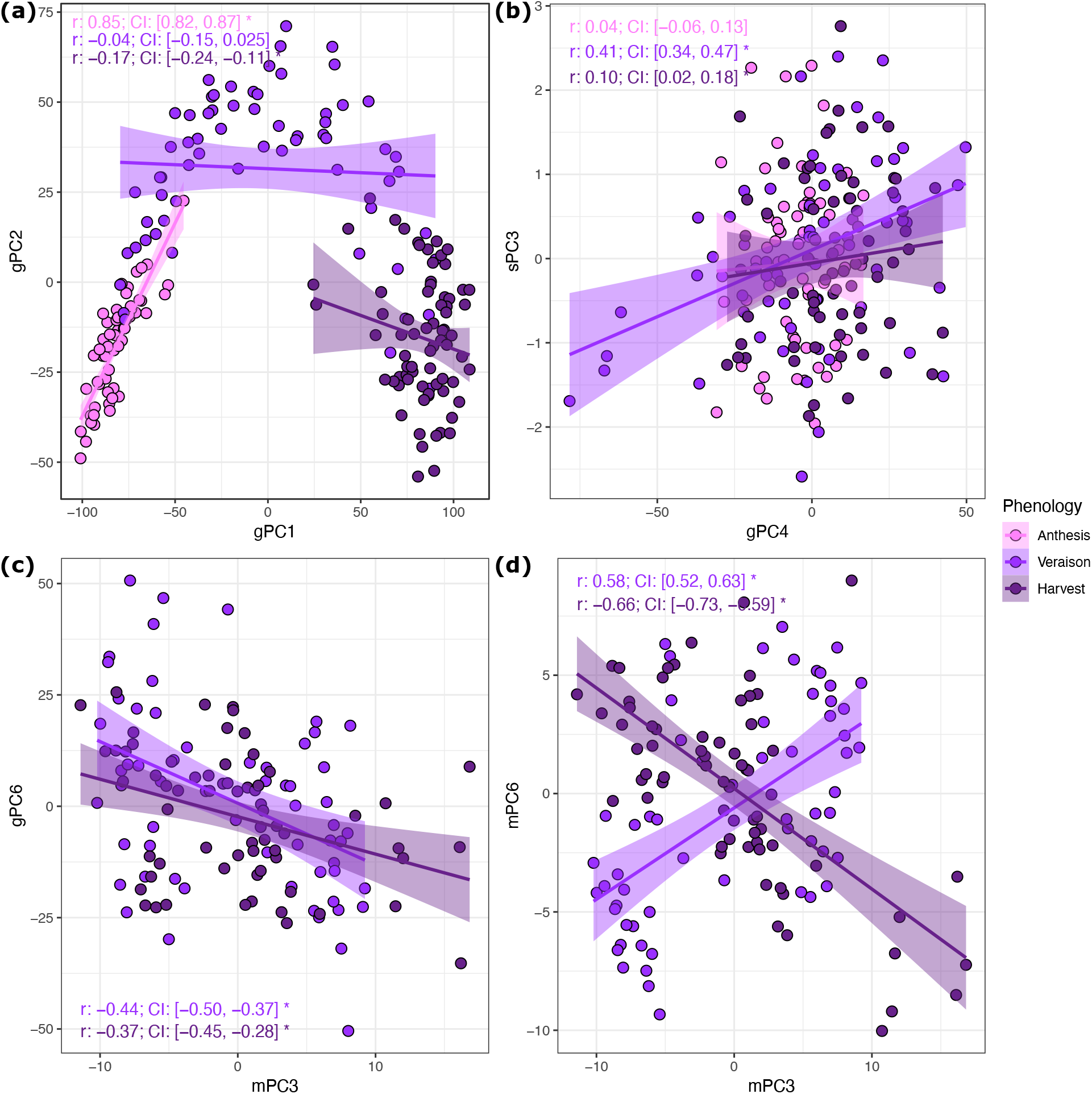
Example correlations within and between data modalities over the course of the season. (A) Example correlation showing a strong within-modality correlation between the ionomics gPC1 and gPC2 at anthesis. Pearson correlations by phenological stage and CIs derived from 10000 random 90% draws are shown for each panel. Generally speaking, CIs overlapping with 0 were not accepted as significant. (B) Example correlation showing one of the stronger between-modality correlations between the gene expression gPC4 and morphology (shape) sPC3 at veraison. (C) Example correlation of a relationship that is present multiple times over the course of the season between metabolomics mPC3 and gene expression gPC6 at both veraison and harvest. (D) Example correlation that is dynamic over the course of the growing season between the ionomics mPC3 and mPC6.

## Data Availability

Ionomics data are available at 10.6084/m9.figshare.13200980. Metabolomics data are available at 10.6084/m9.figshare.13201043. Gene expression data are available in the Sequence Read Archive under BioProject PRJNA674915. Leaf scans and leaf landmarks are available at 10.6084/m9.figshare.13200953. Weather and physiology data are available at 10.6084/m9.figshare.13198682 and 10.6084/m9.figshare.13201016, respectively.

## Code Availability

All code for this paper including shell scripts for RNAseq analysis and Jupyter Notebooks for data analysis in R can be found on the Vitis Underground GitHub (https://github.com/PGRP1546869/mt_vernon_2017_leaf).

## Author Contributions

AJM, DHC, AF, LGK, MK, JPL, and QM designed the experiment. ZNH, LLK, MA, JFS, ZM, NB, EF, and JPL contributed to sample collection and sample processing. ZNH, LLK, JFS, and MA contributed to data analysis. ZNH and AJM contributed to the writing of the manuscript. All authors contributed to manuscript editing.

## Acknowledgments

This work was funded by the National Science Foundation Plant Genome Research Project 1546869. We thank members of the Miller lab at Saint Louis University and the Donald Danforth Plant Science Center, members of the Kovacs Lab at Missouri State University, members of the Kwasniewski Lab at the University of Missouri, and members Londo Lab at the USDA-ARS Grape Research Unit for vineyard sampling and sample processing. We express special thanks to Matthew Rubin and Elizabeth Kellogg at the Donald Danforth Plant Science Center for statistical wisdom and valuable comments on the manuscript.

## Notes

### Competing Interest Statement

The authors have declared no competing interest.

## References

Anders S, Huber W. 2010. Differential expression analysis for sequence count data. Genome biology 11: R106.

Anders S, Pyl PT, Huber W. 2010. HTSeq: Analysing high-throughput sequencing data with Python.

Anesi A, Stocchero M, Dal Santo S, Commisso M, Zenoni S, Ceoldo S, Tornielli GB, Siebert TE, Herderich M, Pezzotti M, et al. 2015. Towards a scientific interpretation of the terroir concept: plasticity of the grape berry metabolome. BMC plant biology 15: 191.

Bavaresco L, Lovisolo C. 2015. Effect of grafting on grapevine chlorosis and hydraulic conductivity. VITIS-Journal of Grapevine Research.

Baxter I. 2010. Ionomics: The functional genomics of elements. Briefings in functional genomics 9: 149–156.

Baxter IR, Gustin JL, Settles AM, Hoekenga OA. 2013. Ionomic characterization of maize kernels in the intermated B73× Mo17 population. Crop science 53: 208–220.

Berdeja M, Nicolas P, Kappel C, Dai ZW, Hilbert G, Peccoux A, Lafontaine M, Ollat N, Gomès E, Delrot S. 2015. Water limitation and rootstock genotype interact to alter grape berry metabolism through transcriptome reprogramming. Horticulture research 2: 15012.

Bolger AM, Lohse M, Usadel B. 2014. Trimmomatic: a flexible trimmer for Illumina sequence data. Bioinformatics 30: 2114–2120.

Bushnell B. 2017. BBTools software package. URL http://sourceforge.net/projects/bbmap.

Canaguier A, Grimplet J, Di Gaspero G, Scalabrin S, Duchêne E, Choisne N, Mohellibi N, Guichard C, Rombauts S, Le Clainche I, et al. 2017. A new version of the grapevine reference genome assembly (12X.v2) and of its annotation (VCost.v3). Genomics data 14: 56–62.

Canas S, Assunção M, Brazão J, Zanol G, Eiras-Dias JE. 2015. Phenolic compounds involved in grafting incompatibility of Vitis spp: development and validation of an analytical method for their quantification. Phytochemical analysis: PCA 26: 1–7.

Chitarra W, Perrone I, Avanzato CG, Minio A, Boccacci P, Santini D, Gilardi G, Siciliano I, Gullino ML, Delledonne M, et al. 2017. Grapevine Grafting: Scion Transcript Profiling and Defense-Related Metabolites Induced by Rootstocks. Frontiers in plant science 8.

Chitwood DH, Klein LL, O’Hanlon R, Chacko S, Greg M, Kitchen C, Miller AJ, Londo JP. 2015. Latent developmental and evolutionary shapes embedded within the grapevine leaf. New Phytologist 210: 343–355.

Chitwood DH, Mullins J, Migicovsky Z, Frank M, VanBuren R, Londo JP. 2020. Vein-to-blade ratio is an allometric indicator of climate-induced changes in grapevine leaf size and shape. bioRxiv: 2020.05.20.106906.

Chitwood DH, Ranjan A, Martinez CC, Headland LR, Thiem T, Kumar R, Covington MF, Hatcher T, Naylor DT, Zimmerman S, et al. 2014. A modern ampelography: a genetic basis for leaf shape and venation patterning in grape. Plant physiology 164: 259–272.

Chitwood DH, Rundell SM, Li DY, Woodford QL, Yu TT, Lopez JR, Greenblatt D, Kang J, Londo JP. 2016. Climate and Developmental Plasticity: Interannual Variability in Grapevine Leaf Morphology. Plant physiology 170: 1480–1491.

Cookson SJ, Ollat N. 2013. Grafting with rootstocks induces extensive transcriptional re-programming in the shoot apical meristem of grapevine. BMC plant biology 13: 147.

Corso M, Vannozzi A, Ziliotto F, Zouine M, Maza E, Nicolato T, Vitulo N, Meggio F, Valle G, Bouzayen M, et al. 2016. Grapevine Rootstocks Differentially Affect the Rate of Ripening and Modulate Auxin-Related Genes in Cabernet Sauvignon Berries. Frontiers in plant science 7: 69.

Csardi G, Nepusz T, Others. 2006. The igraph software package for complex network research. InterJournal, complex systems 1695: 1–9.

Cuadros-Inostroza A, Ruíz-Lara S, González E, Eckardt A, Willmitzer L, Peña-Cortés H. 2016. GC-MS metabolic profiling of Cabernet Sauvignon and Merlot cultivars during grapevine berry development and network analysis reveals a stage- and cultivar-dependent connectivity of primary metabolites. Metabolomics: Official journal of the Metabolomic Society 12: 39.

Dal Santo S, Fasoli M, Negri S, D’Incà E, Vicenzi N, Guzzo F, Tornielli GB, Pezzotti M, Zenoni S. 2016. Plasticity of the Berry Ripening Program in a White Grape Variety. Frontiers in plant science 7: 970.

Dal Santo S, Tornielli GB, Zenoni S, Fasoli M, Farina L, Anesi A, Guzzo F, Delledonne M, Pezzotti M. 2013. The plasticity of the grapevine berry transcriptome. Genome biology 14: r54.

Degu A, Hochberg U, Sikron N, Venturini L, Buson G, Ghan R, Plaschkes I, Batushansky A, Chalifa-Caspi V, Mattivi F, et al. 2014. Metabolite and transcript profiling of berry skin during fruit development elucidates differential regulation between Cabernet Sauvignon and Shiraz cultivars at branching points in the polyphenol pathway. BMC plant biology 14: 188.

Dobin A, Davis CA, Schlesinger F, Drenkow J, Zaleski C, Jha S, Batut P, Chaisson M, Gingeras TR. 2013. STAR: ultrafast universal RNA-seq aligner. Bioinformatics 29: 15–21.

Dryden IL, Mardia KV. 2016. Statistical Shape Analysis: With Applications in R. John Wiley & Sons.

Fabres PJ, Collins C, Cavagnaro TR, Rodríguez López CM. 2017. A Concise Review on Multi-Omics Data Integration for Terroir Analysis in Vitis vinifera. Frontiers in plant science 8: 1065.

Ferlito F, Distefano G, Gentile A, Allegra M, Lakso AN, Nicolosi E. 2020. Scion–rootstock interactions influence the growth and behaviour of the grapevine root system in a heavy clay soil. Australian Journal of Grape and Wine Research 26: 68–78.

Fox J, Friendly M, Weisberg S. 2013. Hypothesis tests for multivariate linear models using the car package. The R journal 5: 39–52.

Galet P. 1979. A Practical Ampelography: Grapevine Identification. Comstock Pub. Associates.

Gautier A, Cookson SJ, Lagalle L, Ollat N, Marguerit E. 2020a. Influence of the three main genetic backgrounds of grapevine rootstocks on petiolar nutrient concentrations of the scion, with a focus on phosphorus. OENO One 54: 1–13.

Gautier A, Cookson SJ, Lagalle L, Ollat N, Marguerit E. 2020b. Influence of the three main genetic backgrounds of grapevine rootstocks on petiolar nutrient concentrations of the scion, with a focus on phosphorus. OENO One 54: 1–13.

Gehan MA, Fahlgren N, Abbasi A, Berry JC, Callen ST, Chavez L, Doust AN, Feldman MJ, Gilbert KB, Hodge JG, et al. 2017. PlantCV v2: Image analysis software for high-throughput plant phenotyping. PeerJ 5: e4088.

Grimes DW, Williams LE. 1990. Irrigation Effects on Plant Water Relations and Productivity of Thompson Seedless Grapevines. Crop science 30: 255.

Harris ZN, Kovacs LG, Londo JP. 2017. RNA-seq-based genome annotation and identification of long-noncoding RNAs in the grapevine cultivar ‘Riesling’. BMC genomics 18: 937.

Houle D, Govindaraju DR, Omholt S. 2010. Phenomics: the next challenge. Nature reviews. Genetics 11: 855–866.

Huang S, Chaudhary K, Garmire LX. 2017. More Is Better: Recent Progress in Multi-Omics Data Integration Methods. Frontiers in genetics 8: 84.

Islam MN, Downey F, Ng CKY. 2011. Comparative analysis of bioactive phytochemicals from Scutellaria baicalensis, Scutellaria lateriflora, Scutellaria racemosa, Scutellaria tomentosa and Scutellaria wrightii by LC-DAD-MS. Metabolomics: Official journal of the Metabolomic Society 7: 446–453.

Jaillon O, Aury J-M, Noel B, Policriti A, Clepet C, Casagrande A, Choisne N, Aubourg S, Vitulo N, Jubin C, et al. 2007. The grapevine genome sequence suggests ancestral hexaploidization in major angiosperm phyla. Nature 449: 463–467.

Klein LL, Caito M, Chapnick C, Kitchen C, O’Hanlon R, Chitwood DH, Miller AJ. 2017. Digital Morphometrics of Two North American Grapevines (Vitis: Vitaceae) Quantifies Leaf Variation between Species, within Species, and among Individuals. Frontiers in plant science 8: 373.

Kuhn M. 2013. Predictive Modeling with R and the caret Package. Google Scholar.

Lecourt J, Lauvergeat V, Ollat N, Vivin P, Cookson SJ. 2015. Shoot and root ionome responses to nitrate supply in grafted grapevines are rootstock genotype dependent: Rootstock and nitrogen supply affect grapevine ionome. Australian journal of grape and wine research 21: 311–318.

Lenth R, Singmann H, Love J, Others. 2018. Emmeans: Estimated marginal means, aka least-squares means. R package version 1.

Liaw A, Wiener M, Others. 2002. Classification and regression by randomForest. R news 2: 18–22.

López-Orenes A, Bueso MC, Conesa H, Calderón AA, Ferrer MA. 2018. Seasonal ionomic and metabolic changes in Aleppo pines growing on mine tailings under Mediterranean semi-arid climate. The Science of the total environment 637-638: 625–635.

Love MI, Huber W, Anders S. 2014. Moderated estimation of fold change and dispersion for RNA-seq data with DESeq2. Genome biology 15: 550.

Marasco R, Rolli E, Fusi M, Michoud G, Daffonchio D. 2018. Grapevine rootstocks shape underground bacterial microbiome and networking but not potential functionality. Microbiome 6: 3.

Migicovsky Z, Harris ZN, Klein LL, Li M, McDermaid A, Chitwood DH, Fennell A, Kovacs LG, Kwasniewski M, Londo JP, et al. 2019a. Rootstock effects on scion phenotypes in a ‘Chambourcin’ experimental vineyard. Horticulture Research 6: 1–13.

Mudge K, Janick J, Scofield S, Goldschmidt EE. 2009. A History of Grafting. In: Janick J, ed. CIBA-GEIGY Agrochemicals Tech. Monog. Horticultural Reviews. Hoboken, NJ, USA: John Wiley & Sons, Inc., 437–493.

Mullins MG, Bouquet A, Williams LE. 1992. Biology of the Grapevine. Cambridge University Press.

Oliver SG, Winson MK, Kell DB, Baganz F. 1998. Systematic functional analysis of the yeast genome. Trends in biotechnology 16: 373–378.

Palumbo MC, Zenoni S, Fasoli M, Massonnet M, Farina L, Castiglione F, Pezzotti M, Paci P. 2014. Integrated network analysis identifies fight-club nodes as a class of hubs encompassing key putative switch genes that induce major transcriptome reprogramming during grapevine development. The Plant cell 26: 4617–4635.

Pauli D, Ziegler G, Ren M, Jenks MA, Hunsaker DJ, Zhang M, Baxter I, Gore MA. 2018. Multivariate Analysis of the Cotton Seed Ionome Reveals a Shared Genetic Architecture. G3 8: 1147–1160.

Pouget R. 1990. Histoire de la lutte contre le phylloxéra de la vigne en France: 1868-1895. Histoire des sciences medicales.

Prodhomme D, Valls Fonayet J, Hévin C, Franc C, Hilbert G, de Revel G, Richard T, Ollat N, Cookson SJ. 2019. Metabolite profiling during graft union formation reveals the reprogramming of primary metabolism and the induction of stilbene synthesis at the graft interface in grapevine. BMC plant biology 19: 599.

R Core Team. 2013. R: A language and environment for statistical computing.

Ripley BD. 2002. Modern applied statistics with S. Springer.

Salt DE, Baxter I, Lahner B. 2008. Ionomics and the study of the plant ionome. Annual review of plant biology 59: 709–733.

Savoi S, Wong DCJ, Arapitsas P, Miculan M, Bucchetti B, Peterlunger E, Fait A, Mattivi F, Castellarin SD. 2016. Transcriptome and metabolite profiling reveals that prolonged drought modulates the phenylpropanoid and terpenoid pathway in white grapes (Vitis vinifera L.). BMC plant biology 16: 67.

Savoi S, Wong DCJ, Degu A, Herrera JC, Bucchetti B, Peterlunger E, Fait A, Mattivi F, Castellarin SD. 2017. Multi-Omics and Integrated Network Analyses Reveal New Insights into the Systems Relationships between Metabolites, Structural Genes, and Transcriptional Regulators in Developing Grape Berries (Vitis vinifera L.) Exposed to Water Deficit. Frontiers in plant science 8: 1124.

Soulé M. 1967. PHENETICS OF NATURAL POPULATIONS I. PHENETIC RELATIONSHIPS OF INSULAR POPULATIONS OF THE SIDE-BLOTCHED LIZARD. Evolution; international journal of organic evolution 21: 584–591.

Stein-O’Brien GL, Arora R, Culhane AC, Favorov AV, Garmire LX, Greene CS, Goff LA, Li Y, Ngom A, Ochs MF, et al. 2018. Enter the Matrix: Factorization Uncovers Knowledge from Omics. Trends in genetics: TIG 34: 790–805.

Swift JF, Hall ME, Harris ZN, Kwasniewki MT, Miller AJ. 2020. Grapevine microbiota reflect diversity among compartments and complex interactions within and among root and shoot systems. BioRxiv.

Tandonnet S, Torres TT. 2017. Traditional versus 3’ RNA-seq in a non-model species. Genomics data 11: 9–16.

Tautenhahn R, Patti GJ, Rinehart D, Siuzdak G. 2012. XCMS Online: a web-based platform to process untargeted metabolomic data. Analytical chemistry 84: 5035–5039.

Team RC, Others. 2013. R foundation for statistical computing. Vienna, Austria 3.

Thomas CL, Alcock TD, Graham NS, Hayden R, Matterson S, Wilson L, Young SD, Dupuy LX, White PJ, Hammond JP, et al. 2016. Root morphology and seed and leaf ionomic traits in a Brassica napus L. diversity panel show wide phenotypic variation and are characteristic of crop habit. BMC plant biology 16: 214.

Tramontini S, Vitali M, Centioni L, Schubert A, Lovisolo C. 2013. Rootstock control of scion response to water stress in grapevine. Environmental and Experimental Botany 93: 20–26.

Tweeddale H, Notley-McRobb L, Ferenci T. 1998. Effect of slow growth on metabolism of Escherichia coli, as revealed by global metabolite pool (‘metabolome’) analysis. Journal of bacteriology 180: 5109–5116.

Ubbens J, Cieslak M, Prusinkiewicz P, Stavness I. 2020. Latent Space Phenotyping: Automatic Image-Based Phenotyping for Treatment Studies.

Ubbens JR, Stavness I. 2017. Deep Plant Phenomics: A Deep Learning Platform for Complex Plant Phenotyping Tasks. Frontiers in plant science 8: 1190.

Vitulo N, Forcato C, Carpinelli EC, Telatin A, Campagna D, D’Angelo M, Zimbello R, Corso M, Vannozzi A, Bonghi C, et al. 2014. A deep survey of alternative splicing in grape reveals changes in the splicing machinery related to tissue, stress condition and genotype. BMC plant biology 14: 99.

Walker MA, Lund K, Agüero C, Riaz S, Fort K, Heinitz C, Romero N. 2014. BREEDING GRAPE ROOTSTOCKS FOR RESISTANCE TO PHYLLOXERA AND NEMATODES - IT’S NOT ALWAYS EASY. Acta Horticulturae: 89–97.

Warschefsky EJ, Klein LL, Frank MH, Chitwood DH, Londo JP, von Wettberg EJB, Miller AJ. 2016. Rootstocks: Diversity, Domestication, and Impacts on Shoot Phenotypes. Trends in plant science 21: 418–437.

Wickham H. 2016. ggplot2: Elegant Graphics for Data Analysis. Springer.

Williams BR, Edwards CE, Kwasniewski MT, Miller AJ. 2020. Epigenomic patterns reflect irrigation and grafting in the grapevine clone ‘Chambourcin’. bioRxiv.

Williams LE, Grimes DW. 1987. Modelling vine growth-development of a data set for a water balance subroutine. In: Proceedings of the Sixth Australian Wine Industry Technical Conference. 169–174.

Wong DCJ, Matus JT. 2017. Constructing Integrated Networks for Identifying New Secondary Metabolic Pathway Regulators in Grapevine: Recent Applications and Future Opportunities. Frontiers in plant science 8: 505.

Zamboni A, Di Carli M, Guzzo F, Stocchero M, Zenoni S, Ferrarini A, Tononi P, Toffali K, Desiderio A, Lilley KS, et al. 2010. Identification of putative stage-specific grapevine berry biomarkers and omics data integration into networks. Plant physiology 154: 1439–1459.

Ziegler G, Terauchi A, Becker A, Armstrong P, Hudson K, Baxter I. 2013. Ionomic Screening of Field-Grown Soybean Identifies Mutants with Altered Seed Elemental Composition. The Plant Genome 6: lantgenome2012.07.0012.

Zombardo A, Crosatti C, Bagnaresi P, Bassolino L, Reshef N, Puccioni S, Faccioli P, Tafuri A, Delledonne M, Fait A, et al. 2020. Transcriptomic and biochemical investigations support the role of rootstock-scion interaction in grapevine berry quality. BMC genomics 21: 468.

