## Supplemental Note 2 for "Root system influence on high dimensional leaf phenotypes over the grapevine growing season"

**Expanded Metabolomics Methods:**

The metabolome represents a catalogue of small molecules present in a tissue, likely stemming from metabolic processes (Oliver *et al.*, 1998; Tweeddale *et al.*, 1998). Metabolomic analysis was completed at veraison and harvest on the 72-vine set. Three mature leaves were sampled from the middle of a randomly selected shoot and immediately flash frozen in liquid nitrogen to prevent any enzymatic reactions and to capture the metabolic state of the leaves when attached to the vine. The frozen leaves were transported to the University of Missouri Enology lab on dry ice and stored at -80˚C until analyzed. Whole leaves were manually ground in liquid nitrogen with a mortar and pestle, 0.5g of powder was weighed into a centrifuge tube, 1.5ml of 1:1 MeOH: ACN was added. Samples were vortexed to suspend leaf particles and sonicated for 20 minutes in an ice bath. Following extraction, samples were centrifuged for 10 minutes at 3,000 g and filtered with a 0.22 PTFE syringe filter into a 1.5ml sample vial before injecting into a Waters XEVO™ QToF LCMS system (Waters Corporation, Milford, MA, USA). Chromatographic separation was achieved using a Waters Acquity ™ Ultra Performance LC H-Class system (Waters Corporation, Milford, MA, USA) equipped with Waters Acquity BEH C18 column (2.1mmx150mm and 1.7um particle size) and a diode array detector. Samples were injected in random order across the sampling periods. The injection volume was set at 2.5ul and the flow rate was set at 0.4 ml/min. The mobile phase consisted of 0.1% formic acid in water (solvent A) and 0.1% formic acid and 5% water in acetaldehyde (solvent B) and the gradient was as follows: 100% A for 0.5 min; 0.5-18min increased to 99% B; 18-19 min. held at 99% B; mobile phase was re-equilibrated for 2 min between runs. Diode array was monitored at 225-500nm. Mass spectrometry was performed on a Xevo™ QTof (Waters Corporation, Milford, MA, USA). The electrospray ionization (ESI) was operated in both positive or negative ionization modes in separate runs. The scan range was set as m/z 50-1500 with 0.2 sec accumulation time. MS settings were as follows: capillary voltage was 2.5kV; cone voltage ramped from 20-40V; collision energy was set to 6V; detector voltage was set to 1950V; desolvation gas was set to 1000 L/hour; cone gas was set to 50 L/hr; source temperature was 120˚C and desolvation temperature was set at 550˚C.

Following analysis, instrument files were converted to .cdf format and uploaded to XCMS online (Tautenhahn *et al.*, 2012) for chromatogram normalization and feature detection via "single job" parameters. Metabolomic features identified as significantly different from background were used as the basis of a principal components (PC) analysis. The top 20 PCs were treated as distinct phenotypes to model according to the experimental design. In PCs that varied significantly by rootstock, features that loaded more than 1.96 standard deviations above or below the mean were fit independently with the same model design.

**References:**

**Oliver SG, Winson MK, Kell DB, Baganz F**. **1998**. Systematic functional analysis of the yeast genome. *Trends in biotechnology* **16**: 373–378.

**Tautenhahn R, Patti GJ, Rinehart D, Siuzdak G**. **2012**. XCMS Online: a web-based platform to process untargeted metabolomic data. *Analytical chemistry* **84**: 5035–5039.

**Tweeddale H, Notley-McRobb L, Ferenci T**. **1998**. Effect of slow growth on metabolism of Escherichia coli, as revealed by global metabolite pool ('metabolome') analysis. *Journal of bacteriology* **180**: 5109–5116.
