## Supplemental Note 1 for "Root system influence on high dimensional leaf phenotypes over the grapevine growing season"

**On the irrigation treatment**

The vineyard in Mount Vernon, MO features a block design randomized over irrigation treatment and the rootstock genotype to which a common scion (Chambourcin) is grafted (or ungrafted in the case of the control). The irrigation treatment comprises three experimental groups: full (replacing 100% of losses from evapotranspiration (ET)), reduced deficit irrigation (RDI; replacing approximately half of ET losses), and none (replacing 0% of ET losses). Estimated ET losses are calculated from an on-site weather station which measures daily precipitation, daily min/max temperature, wind speed, humidity, pressure, and total solar radiance, among others. These values compose an estimated short crop (reference) ET loss (Allen *et al.*, 1998)). Irrigation in the vineyard is then determined by combining that estimate, a grapevine-specific crop coefficient, and the efficiency of the irrigation system. The grapevine-specific crop coefficient (Kc) is the subject of much research (Doorenbos, 1977; Evans *et al.*, 1993; Williams *et al.*, 2003; Campos *et al.*, 2010; Marras *et al.*, 2016; Munitz *et al.*, 2019), but can be generalized to a polynomial function calculated by Grimes and Williams (Grimes & Williams, 1990), and mid-season maximum Kcs have been identified as ranging from 0.6 (describe) to 1.08 (describe). In this Note, we examine the impact of irrigation treatment in 2017 and develop a framework for handling irrigation in our statistical models.

We first examined three measures that should be associated with our irrigation regime assuming the vines were receiving variable amounts of water and/or were experiencing drought stress: 1) soil moisture, 2) water potential, and 3) stomatal conductance. We modelled each trait using the following model factors: Block, Phenology, Irrigation, Rootstock and all possible pairwise combinations (snTable 1). In cases where a trait showed a significant difference as modulated by irrigation (or an interaction with irrigation), we computed post-hoc mean comparisons. In short, we note that soil moisture is variable at veraison between the full and none irrigation treatments (t.ratio=4.03, padj=0.023). However this difference is not shown in comparisons of traits that potentially reflect water stress. In these cases, there is a difference in water potential at veraison between the RDI and none treatments (t.ratio=4.19, padj=0.013), and there is no difference based on irrigation in stomatal conductance. Despite the observed differences in water potential, the differences do not seem to be so severe that the irrigation treatment induced drought stress (snFig. 1). Van Leeuwen et al. (Van Leeuwen *et al.*, 2009) reported ranges of water potential indicative of stress in two categories: weak to moderate water deficit (-9 to -13 bar) and moderate to severe stress (-13 to -14 bar). Additionally, Flexas et al. (Flexas *et al.*, 2006) reported stomatal conductance ranges indicative of stress including moderate stress ( $< 0.15$ mol/m$^2$/s) and severe stress ($< 0.05$ mol/m$^2$/s). While we certainly observed instances of water stress based on both of these categorizations, they do not seem to reflect the irrigation treatment.

snTable 1.

| | Model Factor | SS | df | F | p |
|---|---|---|---|---|---|
| Soil Moisture | Irrigation | 3.705298 | 2 | 0.245568 | 0.7825 |
| Soil Moisture | Block | 187.279664 | 2 | 12.4119301 | **9.06E-06** |
| Soil Moisture | Rootstock | 9.357492 | 3 | 0.4134442 | 0.7436 |
| Soil Moisture | Phenology | 4282.033527 | 2 | 283.7910935 | **8.48E-56** |
| Soil Moisture | Irrigation:Block | 40.437338 | 4 | 1.3399891 | 0.2570 |
| Soil Moisture | Irrigation:Rootstock | 65.420995 | 6 | 1.4452554 | 0.1999 |
| Soil Moisture | **Irrigation:Phenology** | 103.289907 | 4 | 3.4227611 | **0.0101** |
| Soil Moisture | Block:Rootstock | 40.368356 | 6 | 0.8918021 | 0.5022 |
| Soil Moisture | Block:Phenology | 200.998241 | 4 | 6.6605633 | **5.15E-05** |
| Soil Moisture | Rootstock:Phenology | 19.176968 | 6 | 0.4236502 | 0.8626 |
| Soil Moisture | Residuals | 1327.804005 | 176 | | |
| Stomatal Conductance | Irrigation | 0.014462271 | 2 | 1.8917976 | 0.1539 |
| Stomatal Conductance | Block | 0.003624752 | 2 | 0.4741508 | 0.6232 |
| Stomatal Conductance | Rootstock | 0.01114674 | 3 | 0.9720639 | 0.4072 |
| Stomatal Conductance | Phenology | 0.062432311 | 2 | 8.1667187 | **0.0004** |
| Stomatal Conductance | Irrigation:Block | 0.022383646 | 4 | 1.463993 | 0.2152 |
| Stomatal Conductance | Irrigation:Rootstock | 0.024047769 | 6 | 1.0485562 | 0.3957 |
| Stomatal Conductance | Irrigation:Phenology | 0.005697191 | 4 | 0.3726224 | 0.8279 |
| Stomatal Conductance | Block:Rootstock | 0.033476007 | 6 | 1.4596562 | 0.1947 |
| Stomatal Conductance | Block:Phenology | 0.079669605 | 4 | 5.2107574 | **0.0005** |
| Stomatal Conductance | Rootstock:Phenology | 0.081232411 | 6 | 3.5419813 | **0.0025** |
| Stomatal Conductance | Residuals | 0.66891336 | 175 | | |
| Water Potential | Irrigation | 29.537347 | 2 | 4.1984862 | **0.0165** |
| Water Potential | Block | 16.747156 | 2 | 2.3804678 | 0.0955 |
| Water Potential | Rootstock | 7.169907 | 3 | 0.6794282 | 0.5657 |
| Water Potential | Phenology | 102.149803 | 2 | 14.5197382 | **1.46E-06** |
| Water Potential | Irrigation:Block | 19.000741 | 4 | 1.350398 | 0.2532 |
| Water Potential | Irrigation:Rootstock | 22.971551 | 6 | 1.0884045 | 0.3711 |
| Water Potential | **Irrigation:Phenology** | 64.004491 | 4 | 4.5488509 | **0.0016** |
| Water Potential | Block:Rootstock | 26.274745 | 6 | 1.2449116 | 0.2856 |
| Water Potential | Block:Phenology | 87.96463 | 4 | 6.2517173 | **0.0001** |
| Water Potential | Rootstock:Phenology | 10.490926 | 6 | 0.4970657 | 0.8100 |
| Water Potential | Residuals | 619.10088 | 176 | | |

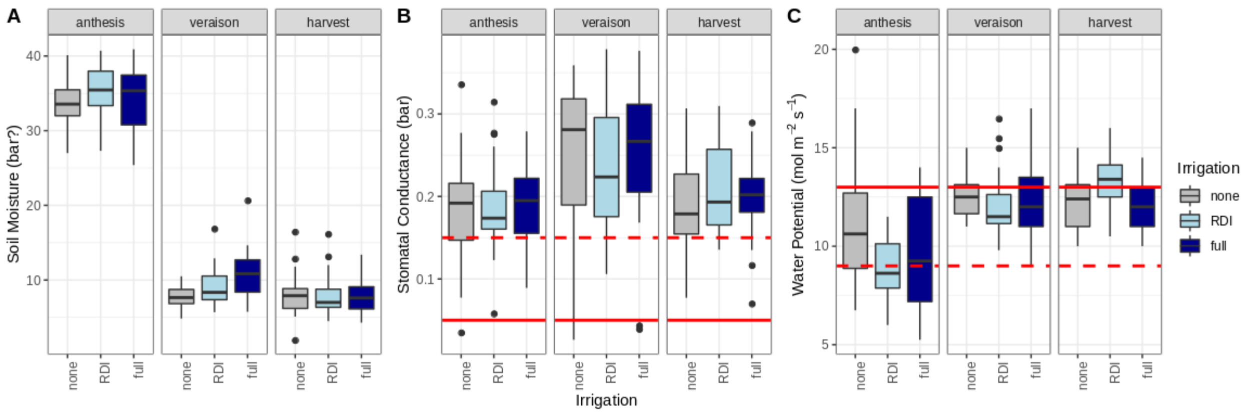


snFig 1. Measured traits expected to respond to drought treatments. A) Soil moisture shown across the season and averaged across irrigation treatments. B) Stomatal conductance of leaves. The horizontal red lines indicate the maximal values for severe stress threshold (solid) and the moderate stress threshold (dashed) as reported by Flexas et. al (2006). C) Leaf water potential with horizontal red lines indicating the minimal values for moderate-to-severe stress (solid) and weak-to-moderate stress (dashed)

We then sought to understand why the irrigation treatment was not producing the anticipated stress responses. To do this, we downloaded four years of weather data from the on-site weather station comprising 2015 (the first complete year for which data exist) through 2018 (the last complete year for which data existed). For each year, we looked at how the cumulative rainfall was associated with the predicted estimates of grapevine evapotranspiration losses over a range of values derived from the literature (sources from Jupyter). In this analysis, we used the polynomial function derived by Grimes and Williams (1990) with adjusted y-intercepts to account for the range of literature derived values. Daily predicted ET losses were modified by multiplying those values by daily crop coefficient estimates. For each year but 2016, rainfall overshadowed the predicted ET losses across the entire year (snFig 2a). In 2017, we note a rapid accumulation of rainfall immediately prior to sampling at anthesis, a lack of rainfall leading up to sampling at veraison, and slightly longer lack of rainfall leading up to harvest.

Finally, we focused on the 2017 sampling window to see if sampling dates were prefaced by significant rainfall events, potentially “washing away” the effects of the irrigation treatments (snFig 2b). At anthesis, five of the ten days leading up to sampling experienced rainfall, three of which more than enough to fully replace evapotranspiration losses. Leading up to veraison, two days experienced rainfall following the second longest period of no rainfall in the sampling window. Leading to harvest, there was only one day of rainfall following the longest period of the sampling window without rainfall. Given that grapevines can typically recover from prolonged drought stress in a matter of days (Hochberg *et al.*, 2017), we propose that the vines were experiencing minimum stress throughout the season, with effects likely being apparent only during windows of decreased rainfall. However, both the anthesis and harvest sampling dates were sufficiently after rainfall events for the vines to have recovered or to have at least begun recovering from the drought and irrigation deficits. At veraison, there does not seem to have been enough time to overcome the prolonged dry period as evidenced by soil moisture, but the plants still do not exhibit signatures of stress as measured through water potential or stomatal conductance. It is possible, though, that while the vines are experiencing a lack of water, they are influenced by previous irrigation treatments, either from previous seasons, previous periods of prolonged rainfall reduction, or a combination of both. This is supported by our previous work where we showed that for a single pre-veraison, pre-irrigation treatment time point, the vines still showed signatures of the irrigation treatment in leaf shape, and leaf ion concentrations (Migicovsky *et al.*, 2019). This sampling event followed a short period of rainfall reduction and three years over which there were likely periods of decreased rainfall (like the window following ~harvest in 2015). Collectively, we have opted to only minimally include irrigation in the statistical models of this analysis. For each model fit, we included irrigation as a single, non-interacting fixed effect under the assumption that irrigation/ stress will have a minimal impact on the results of this study.

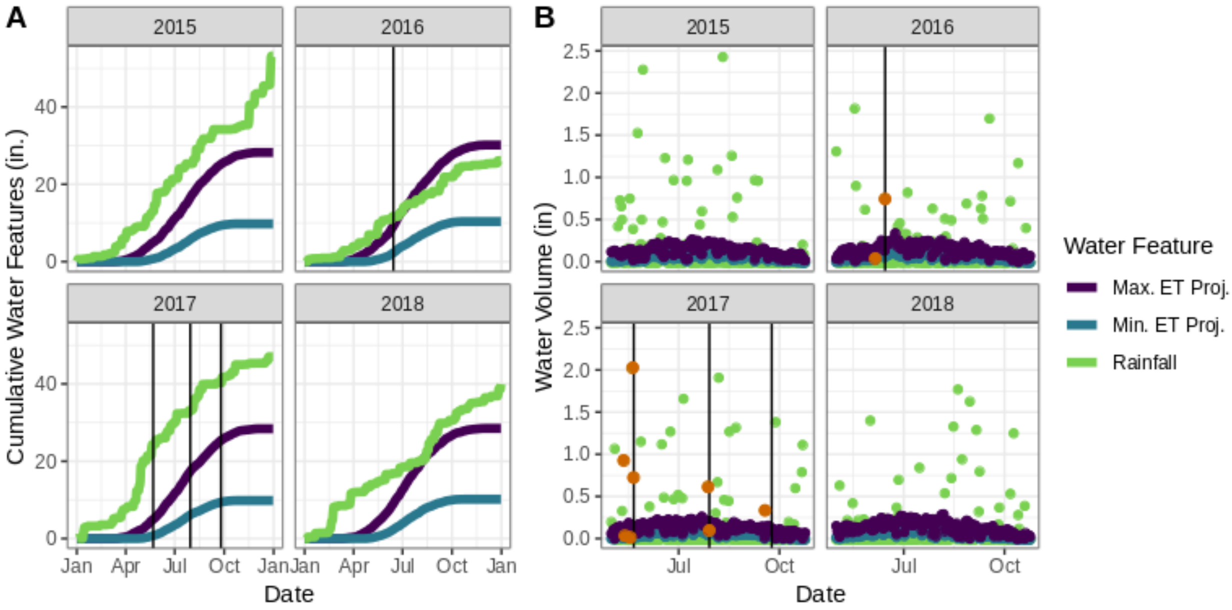


snFig 2. Rainfall as compared to projected ET losses at two scales. A) Cumulative rainfall as compared to cumulative ET loss projections from a modified form of Grimes and Williams estimated for all relevant years in which there was complete data. Dark vertical lines indicate sampling events for 2016 and 2017. B) Daily rainfall and predicted ET losses over an extended sampling window. Dark vertical lines indicate sampling events in 2017, and dark orange points indicate rainfall events up to ten days preceding vineyard sampling.

References:

**Allen RG, Pereira LS, Raes D, Smith M. 1998**. FAO Irrigation and drainage paper No. 56. *Rome: Food and Agriculture Organization of the United Nations* **56**: e156.

**Campos I, Neale CMU, Calera A, Balbontín C, González-Piqueras J. 2010**. Assessing satellite-based basal crop coefficients for irrigated grapes (Vitis vinifera L.). *Agricultural water management* **98**: 45–54.

**Doorenbos J**. Pruitt., WO (1977). Guidelines for predicting crop water requirements. FAO Irrigation and Drainage Paper 24. *Food and Agriculture Organization. Rome.*

**Evans RG, Spayd SE, Wample RL, Kroeger MW, Mahan MO**. **1993**. Water use of Vitis vinifera grapes in Washington. *Agricultural water management* **23**: 109–124.

**Flexas J, Bota J, Galmés J, Medrano H, Ribas-Carbó M**. **2006**. Keeping a positive carbon balance under adverse conditions: responses of photosynthesis and respiration to water stress. *Physiologia plantarum* **127**: 343–352.

**Grimes DW, Williams LE**. **1990**. Irrigation Effects on Plant Water Relations and Productivity of Thompson Seedless Grapevines. *Crop science* **30**: 255.

**Hochberg U, Bonel AG, David-Schwartz R, Degu A, Fait A, Cochard H, Peterlunger E, Herrera JC**. **2017**. Grapevine acclimation to water deficit: the adjustment of stomatal and hydraulic conductance differs from petiole embolism vulnerability. *Planta* **245**: 1091–1104.

**Marras S, Achenza F, Snyder RL, Duce P, Spano D, Sirca C**. **2016**. Using energy balance data for assessing evapotranspiration and crop coefficients in a Mediterranean vineyard. *Irrigation Science* **34**: 397–408.

**Migicovsky Z, Harris ZN, Klein LL, Li M, McDermaid A, Chitwood DH, Fennell A, Kovacs LG, Kwasniewski M, Londo JP, *et al.* 2019**. Rootstock effects on scion phenotypes in a ‘Chambourcin’ experimental vineyard. *Horticulture research* **6**: 64.

**Munitz S, Schwartz A, Netzer Y**. **2019**. Water consumption, crop coefficient and leaf area relations of a Vitis vinifera cv. ‘Cabernet Sauvignon’ vineyard. *Agricultural water management* **219**: 86–94.

**Van Leeuwen C, Trégoat O, Choné X, Bois B, Pernet D, Gaudillère J-P**. **2009**. Vine water status is a key factor in grape ripening and vintage quality for red Bordeaux wine. How can it be assessed for vineyard management purposes? *OENO One* **43**: 121–134.

**Williams LE, Phene CJ, Grimes DW, Trout TJ**. **2003**. Water use of mature Thompson Seedless grapevines in California. *Irrigation Science* **22**: 11–18.
